# Leveraging biological complexity to predict patch occupancy in a recent host range expansion

**DOI:** 10.1101/2020.04.29.069559

**Authors:** M. L. Forister, C. S. Philbin, Z. H. Marion, C. A. Buerkle, C. D. Dodson, J. A. Fordyce, G. W. Forister, S. L. Lebeis, L. K. Lucas, C. C. Nice, Z. Gompert

## Abstract

Specialized plant-insect interactions are a defining feature of life on earth, yet we are only beginning to understand the factors that set limits on host ranges in herbivorous insects. To understand the colonization of alfalfa by the Melissa blue butterfly, we quantified arthropod assemblages and plant metabolites across a wide geographic region, while controlling for climate and dispersal inferred from population genomic variation. The presence of the butterfly is successfully predicted by direct and indirect effects of plant traits and interactions with other species. Results are consistent with the predictions of a theoretical model of parasite host range in which specialization is an epiphenomenon of the many barriers to be overcome rather than a consequence of trade-offs in developmental physiology.

**One sentence summary:** The formation of a novel plant-insect interaction can be predicted with a combination of biotic and abiotic factors, with comparable importance revealed for metabolomic variation in plants and interactions with mutualists, competitors and enemies.

## Main text

Emerging infectious diseases and crop pests are examples of host range expansion in which an organism with a parasitic life style colonizes and successfully utilizes a novel host (*1*). Many aspects of host range are poorly understood, including why most herbivorous insects and other parasites are specialized and the conditions under which new host-parasite interactions develop and persist (*1*, *2*). Reductionist approaches in focal systems have revealed key aspects of host recognition (*3*) and other relevant mechanisms (*4*), but by design do not encompass context dependence including interacting species and abiotic variation. Ecological studies of host range, in contrast, might quantify context dependence but have not always included modern genomic and metabolomic approaches (*5*). Here we use the colonization of alfalfa, *Medicago sativa*, by the Melissa blue butterfly*, Lycaeides melissa* (Fig. 1) to present what is to our knowledge the most thorough picture of a recent (within the last 200 years) host range expansion in terms of number of populations studied and breadth of interacting species and host traits characterized.

**Fig. 1.**
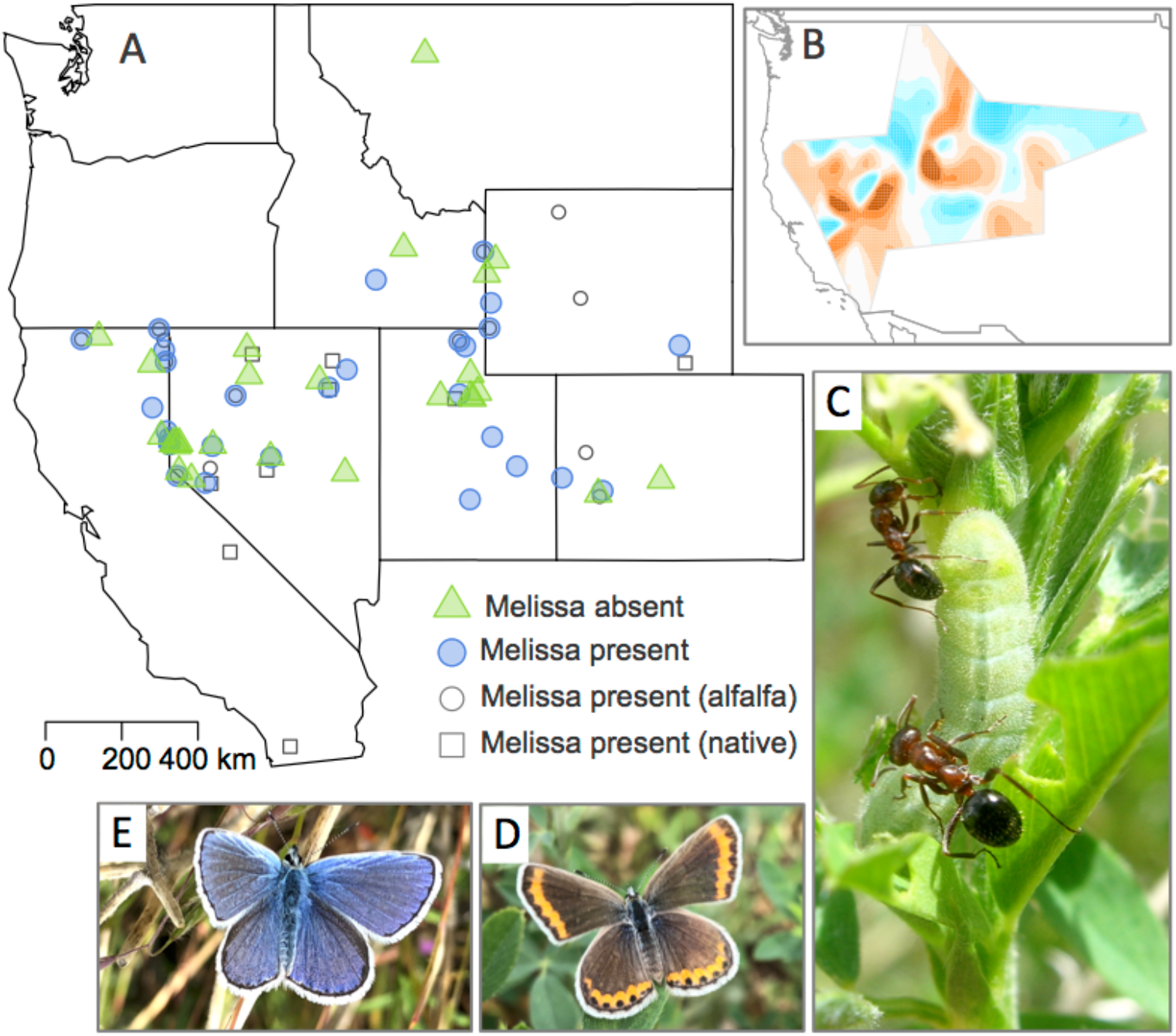
Map of study locations, dispersal surface, and images of butterflies, ants and caterpillar. (**A**) Solid symbols (circles and triangles) are focal alfalfa locations from which arthropods and plants were collected: blue sites are locations where the Melissa blue butterfly (*Lycaeides melissa*) has colonized the novel host; green sites are alfalfa locations not colonized by the butterfly. Open symbols (circles and squares) are locations used in the quantification of gene flow; in some cases (where an open circle appears within a blue circle), sites were represented in both datasets. (**B**) Effective migration surface used to generate covariates representing rates of effective dispersal (blue is faster than average, red is slower). (**C**) *L. melissa* caterpillar being tended by mutualist ants on alfalfa (photo by CCN). (**D**) Female and (**E**) male *L. melissa* butterflies (photos by MLF).

Theoretical work in this area can be divided into two partially-overlapping groups, those that emphasize developmental performance (including trade-offs in the ability to use different hosts), and those that stress opportunity and constraint imposed by exogenous factors, primarily natural enemies (*6*) and geography (*7*). Although developmental trade-offs in host use are rare (*8*), it is clear that plant defenses are a barrier to insect colonization, as performance is often reduced for herbivores in experiments with novel vs ancestral hosts (*9*). What we do not know is whether the magnitude of performance effects studied in the lab will be informative under field conditions. Predation pressure could, for example, remove all opportunity for successful development on a novel host that would otherwise be suitable. Equally unknown is whether variation within and among host populations might have compensatory effects, such that a direct negative effect of a particular toxin on an herbivore is balanced by similar effects on a competitor.

*Lycaeides melissa* is widespread in western North America, where it can be found in association with native legume (Fabaceae) host plants, and typically persists in isolated subpopulations connected by limited gene flow (*10*). The association with alfalfa is heterogenous, and most often occurs in areas where the plant has escaped cultivation. Alfalfa was introduced to western North America in the mid 1800s (*10*), and is a poor food plant for *L. melissa* caterpillars, which develop into adults that can be up to 70% smaller than individuals on native hosts, with direct (*11*) and indirect fitness consequences (*12*). The use of *M. sativa* does not appear to be constrained by genetic, developmental trade-offs in *L. melissa* or a lack of genetic variation in ability to utilize that host (*13*, *14*). Nevertheless, unoccupied patches of *M. sativa* have remained unoccupied by the butterfly for years or even decades, even in close proximity to occupied patches (*15*). We studied that heterogeneity using more than 1,600 individual plants from 56 alfalfa locations with and without *L. melissa* (Fig. 1A). We find that roughly three-quarters of the variation in *L. melissa* presence and absence at the landscape scale can be predicted with a structural equation model (Fig. 2) and a suite of variables that includes metabolomic variation, host patch area, the abundance of interacting arthropods, and dispersal (relative rates of effective migration; Fig. 1B). The success of the model is also apparent in cross-validation (Fig. 2) and null simulations of site-level properties (Supplementary Figure S7).

**Fig. 2.**
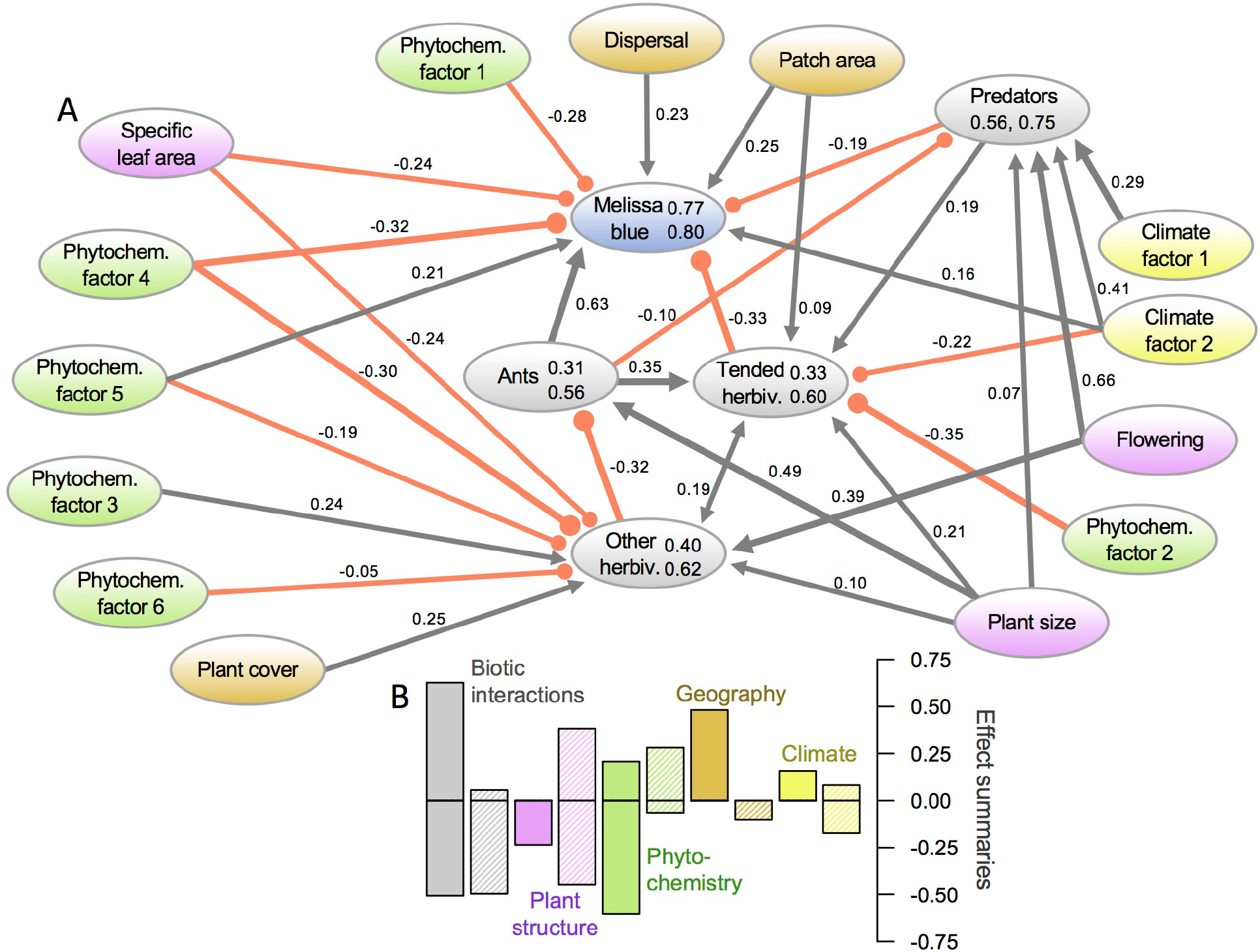
Structural equation model and summary of direct and indirect effects. (**A**) Path diagram illustrates coefficients estimated in structural equation model predicting *L. melissa* presence and absence across the landscape, as well as abundance of ants, tended herbivores, other herbivores and predators (model fit: Fisher’s C = 67.66, P = 0.995). Negative effects are indicated by red lines, positive effects by gray lines; width of lines are scaled to the magnitude of the coefficients. For the endogenous variables, two numbers are shown within ovals: R^2^ values (on top) and observed-vs-predicted correlations (below) from leave-one-out cross validation. Color coding of exogenous variables indicates plant metabolomics data (green), plant structural traits (violet), geographic variables (brown), and climate (yellow); color coding also corresponds to bar chart (**B**), which summarizes relative magnitude of direct and indirect effects (solid and hashed bars, respectively), both positive and negative. For example, climate has a modest positive direct effect, a smaller positive indirect effect (mediated through tended herbivores), and a larger negative indirect effect (through predators).

Like most butterflies in the family Lycaenidae, *L. melissa* caterpillars engage in a facultative mutualism with ants (Fig. 1C), where caterpillars produce specialized secretions in exchange for protection from natural enemies (*16*). Previous experimental work in this system found that excluding ants from individual plants reduced caterpillar survival (*17*). We find here that ant abundance is the most influential variable or control on *L. melissa* presence across the 56 sites (Fig. 2, Fig. 3A). This is true even when considering the fact that ants facilitate hemipterans (aphids, treehoppers, and other myrmecophiles), which in turn have a negative competitive effect on *L. melissa* (Fig. 3B). The balance of ant and hempiteran effects is such that the negative effect of the latter is most influential at intermediate ant densities (Fig. 3D). Similar complexity arises through direct and indirect effects of metabolomic variation. Phytochemical factor 4 has a direct negative association with *L. melissa* presence (Fig. 3C), but an indirect positive effect mediated through other herbivores and their effect on ants (Fig. 2). That axis of plant variation is positively associated with a number of alkaloids, among other compounds, with potential herbivore toxicity (see Supplementary Table S4 and Figure S5).

**Fig. 3.**
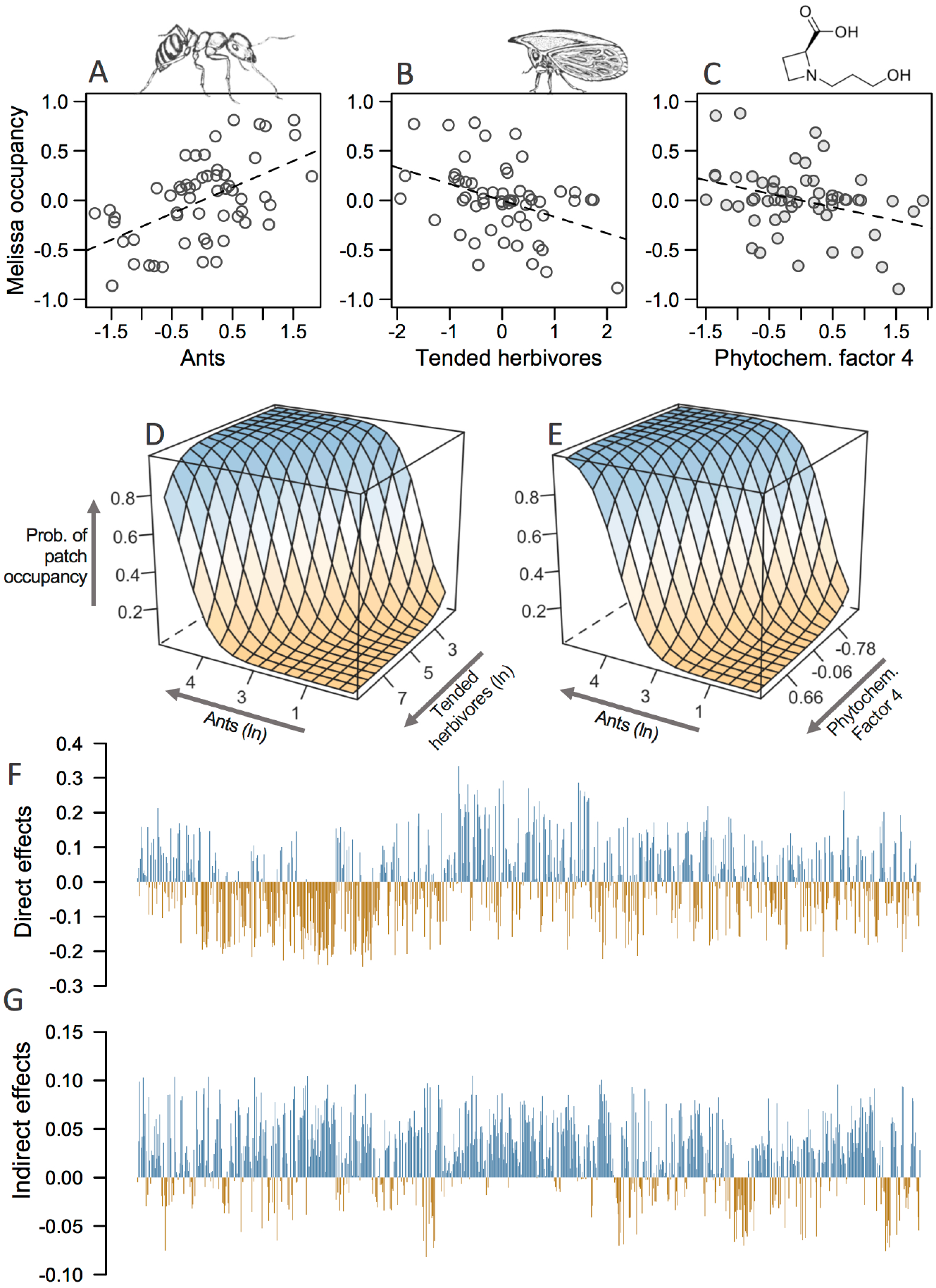
Illustration of effects for a subset of variables predicting *L. melissa* presence and absence across the landscape. (**A**, **B** and **C**) Partial effects of ants, tended herbivores, and phytochemical factor 4 on *L. melissa* occupancy; in other words, these are the effects of those individual factors while controlling for other factors predicting occupancy (see paths in Fig. 2). (**D** and **E**) Predicted probability of patch occupancy across a range of values for ant and tended herbivore abundance (**D**) and for ant abundance and phytochemical factor 4 (**E**) where it can be seen, for example, that at high values of phytochemical factor 4, a higher abundance of ants is needed before the probability of occupancy rises. (**F** and **G**) direct and indirect effects of 849 metabolomic features, both positive (blue) and negative (orange); see text for more details on calculation of individual effects. *Formica* ant and *Campylenchia* treehopper (one of the more abundant ant-tended herbivores at our study sites) illustrations by MLF; the alkaloid shown above (**C**) is medicanine (see Supplementary Figure S5 for additional examples).

Considering the summed totals of direct and indirect effects estimated through path analysis (Fig. 2B), we find that metabolomic variation is associated with the most pronounced, direct negative effects, followed closely (among negative effects) by direct and indirect interactions with other arthropods and then indirect effects of plant structure. The effect of specific leaf area is consistent with a previous experimental study (*18*), but is small compared to both positive and negative indirect effects associated with plant size and the density of flowers mediated through enemies and competitors (Fig. 2B). In terms of positive effects on *L. melissa*, the importance of ants is followed by geographic factors including patch area and dispersal (effective migration rates). These results demonstrate the value of studying plant variation in the context of geography and interacting species. While individual components of the results reported here are consistent with experimental work, other aspects are less accessible to manipulation. Individual metabolites, for example, have a mix of positive and negative direct effects on *L. melissa* (Fig. 3F) as observed in a previous rearing experiment (*18*), while the indirect effects of individual compounds are characterized more by positive effects mediated through numerous other species in the wild (Fig. 3G).

The theory of ecological fitting suggests that novel hosts are colonized if they are "close enough" to native hosts in key traits (*19*–*22*), but we have few cases in which that "close enough" distance has been quantified as we have done in this system. We find diverse factors or controls on colonization that are encountered in multifarious combinations (*23*). When all factors align, butterfly populations persist on the novel host, but the diversity of challenges (plants, enemies, abiotic conditions) undoubtedly makes adaptation to the novel host difficult, especially when some or all of those factors likely shift in character from year to year (*24*). This possibility is consistent with only minimal local adaptation that has been observed in alfalfa-associated populations (*13*). Given these results, we can see the theory of host range evolution approaching maturity: genetic trade-offs are possible, but rare (*25*); instead, it is likely that a balance of factors (both positive and negative) associated with novel host use exist in any system but are only infrequently encountered in combinations that allow host range expansion (*26*–*28*). Thus, generalist herbivores or parasites (with many accumulated hosts) are predictably rare across geographic and phylogenetic scales (*29*, *30*). The complexity of barriers to novel host use and the ecological contingency of colonization challenge our ability to forecast new crop pests or emerging infectious diseases (*1*), but the multi-disciplinary approach illustrated here does raise the promise, at least for herbivorous insects, that expansions of host range can be understood given current technologies and sufficient sampling effort.

## ACKNOWLEDGEMENTS

Thanks to the Hitchcock Center for Chemical Ecology. Thanks to Sarah Flanagan and Kate Bell for help with collections, and Josh Jahner, Su’ad Yoon and Josh Harrison whose travels in the Great Basin discovered many important field sites.

## Author contributions

Overall concept and approach by M.L.F., C.S.P, C.A.B, C.D.D., J.A.F., S.L.L., L.K.K., C.C.N. and Z.G. Data collection by Z.H.M., C.S.P, G.W.F and M.L.F. Data analysis by M.L.F., C.S.P., Z.G., J.A.F., C.C.N., and Z.H.M. All authors read and edited the manuscript.

## Data availability

The data analyzed in this study will be available on Dryad.

## SUPPLEMENTARY MATERIALS

Materials and Methods

Supplementary Results

Figures S1-S7

Tables S1-S13

