## Supplemental material for "Leveraging biological complexity to predict patch occupancy in a recent host range expansion"

*by* Forister, et al.

**This PDF file includes:**

Materials and Methods
Supplementary Results
Supplementary References
Figs. S1 to S7
Tables S1 to S13

*Note: figures are embedded in text, tables are at the end.*

### Materials and Methods

**1. Site identification, plant collections and associated data**

Our goal was to sample a roughly equal number of alfalfa (*Medicago sativa*) locations that had and had not been colonized by the Melissa blue butterfly (*Lycaeides melissa*). Throughout the arid western US, the plant is favored by the butterfly in places where it has escaped cultivation and exists in mostly discrete patches along roadsides and in invaded or partly degraded natural communities. Prior to the field sampling associated with this project in 2017 and 2018, we had accumulated a database of observations of alfalfa locations that support Melissa blue populations, as described in previous publications (*1–3*), as well as (to a lesser extent) a database of alfalfa locations not associated with Melissa presence (*4*). Both presence and absence of the butterfly are relatively stable over time: we have observed presence to be unchanged at locations that we have visited annually for more than 20 years, and focal absences have been tracked for up to 12 years (*4*).

Throughout the summers of 2017 and 2018, we visited a total of 56 sites (Fig. 1A, Supplementary Table S1), mixing presence and absence sites haphazardly in space so as not to confound latitude or longitude with order (date) of sampling or with site status (presence and absence of the butterfly). In other words, sites were visited so that presence and absence sites were interdigitated both in space and time (note that we also include year as a categorical variable in analyses, as described below). In many cases, sites had been previously identified (as mentioned above), while in other cases new sites were discovered during the summers of 2017 and 2018. Because adults have high site fidelity (*5*) and are easily observed, the status (butterfly presence or absence) can in most cases be determined in a single visit. However, *L. melissa* has multiple generations in a season, which makes it possible that a visit could coincide with low density between generations and thus produce a false negative observation (which is more likely early in the summer, as the first and second generations are more discrete, while the second and third tend to overlap). Thus all absence locations were revisited at least once or twice, weeks after the initial sampling date to confirm that a site does not support an *L. melissa* population (the only exception to that was our most northern absence site, in Montana, which was not revisited for logistical reasons). Note that we did not attempt to quantify butterfly abundance, which (unlike presence or absence) would have required multiple visits to reliably estimate.

Following our earlier studies with arthropod communities on alfalfa (*6*, *7*), we used flowering as a guide to phenology, and considered a site appropriate for sampling if at least half of the individual plants were flowering. Some of our previous work with alfalfa arthropods and plant traits has involved repeated sampling at individual locations (*7*), although arthropod communities in alfalfa were found to be relatively stable throughout the summer (*6*), thus the present project utilized single visits to individual locations as a way to maximize effort spent sampling additional sites (Fig. 1A). Most sampling days were in July and August, with fewer in June and September, and an index of flowering was included as a covariate in models (see below). Sampling at a site began with a marker placed in the center of the patch of alfalfa, and 30 individual alfalfa plants were then flagged at random compass directions and distances (up to 25m) from the center of the patch (at a few sites where a patch had less than 30 plants, they were all sampled). In some cases, when patches were essentially linear (for example along a roadside), compass directions were converted to binary orientations forward or backward from the center. The total areal extent (length and width) of the patch was measured and percent cover of alfalfa was visually estimated. Each of the randomly-selected focal plants was measured for size (as a box measurement of length, width and height) and the number of flowering stems was counted. Three mature (but not senescent) leaves were selected haphazardly for leaf toughness measurements using a penetrometer (Chatillon 516 series) through the center of the middle leaflet (*8*).

For metabolomic work (and the measurement of leaf area and mass), three small clusters of leaves (3-5 leaves in each cluster) were collected from the top, middle and lower portions of the plant, thus encompassing as much whole plant phytochemical variation as possible. The three clusters were pooled into a single large coin envelope. Thus, the current study does not quantify intra-individual variation or attempt to separate induced from constitutive defenses (alfalfa is attacked by a wide range of vertebrate and invertebrate herbivores and the vast majority if not all of our sampled plants had been damaged to some extent prior to sampling). All of the envelopes from a single site were stored in an open paper bag for air drying before being delivered to a lab at the University of Nevada, Reno, where drying was completed in a vacuum. After vacuum drying, sample envelopes were kept in plastic bins with desiccating crystals until they were needed for either metabolomic work or area and mass measurements. For the latter (area and mass), five leaves were selected haphazardly from each envelope and weighed to the nearest tenth of a milligram on a microbalance and taped to a sheet of plain white paper that was then scanned. Leaf area (in $cm^2$) was taken using ImageJ software (v. 1.52A) on the scanned images of individual leaves and used to calculate specific leaf area (SLA) as area divided by mass (*8*). Because all of the analyses presented here focus on variation at the patch scale, plant measurements were averaged across plants within a location.

Climatic data for each of the 56 sites was generated as monthly averages for 2008 to 2018 using the *get_prism_monthlys* function from the prism package (*9*) for average daily minimum temperatures, average daily maximum temperatures and precipitations totals (data from the PRISM Climate Group, Oregon State University). These analyses and all others described below (except where noted) were done using the R statistical computing language (*10*).

### 2. Arthropod collections and identification

Arthropods were collected using a sweep net from each of the randomly-selected individual plants at each of the 56 field sites, as we have done previously (*6*, *7*, *11*). Unlike alfalfa in cultivation, alfalfa that has escaped along roadsides and into semi-natural communities tends to grow as larger individuals with open spaces between plants, which facilitates sweep netting of individual plants. Each plant was swept four times in rapid succession, with the collector gradually moving around the plant during sweeping. The collected arthropods were then transferred from the net into a plastic vial with ethanol for storage (an aspirator was used to get most arthropods out of the net, with larger specimens transferred directly into alcohol). We attempted to collect all arthropods large enough to be seen with the naked eye, with the exception of thrips (Thysanoptera) which we have found to be both too small and too numerous on alfalfa for efficient collection. Vials were each labelled by location and plant.

Individual arthropods within samples (vials associated with individual plants) were counted and identified using a dissecting microscope (with 90x maximum magnification) to the lowest possible taxonomic level, which was almost always family and in some cases genus and species, using standard taxonomic keys appropriate to different groups (*12–21*). If a specimen could only be identified to family it was given a morphospecies number that identified unique morphospecies across all of our field sites. This work built on a previous study of alfalfa insects in the Great Basin (*7*), and many of the morphospecies numbers used in the current study were established in that previous work. The only exception to that pipeline involved spiders, which were only counted and not identified to any taxonomic level, such that total spider abundance was used in analyses (as part of the pool of predators at each site).

Following taxonomic identification, each species (or morphospecies) was given one of the following ecological assignments: ant tended herbivores, other herbivores, parasitoids, predators, and ants. These assignments were based partly on our own observations of alfalfa-insect communities, and on literature searches for specific taxa (for example as a way to determine if a particular chewing herbivore could have been feeding on alfalfa or might have more likely been there only as a flower visitor). Ants were treated as their own ecological group for the simple reason that they are considered separately in analyses as mutualists of our focal caterpillars and of other ant tended herbivores. Parasitoids include wasps and flies with some history (based on literature searches) of potentially attacking caterpillars (we also identified a large number of hemipteran parasitoids, but those were not included in these analyses). For analyses reported here, arthropods were totaled within functional (ecological) groups at the plant patch level, and abundances were natural log transformed prior to analyses.

**3. Plant metabolomics**

Individual plants across all sites in each collection year were randomized prior to extraction and analysis. After leaves were vacuum dried and finely ground (using a TissueLyser II, Quiagen; Hilden, Germany), approximately 10 mg of dried tissue was extracted in 2.00 ml of aqueous ethanol (70%), vortexed briefly and sonicated for 15 minutes. The resulting suspension was centrifuged at 500 rpm for 10 min. Aliquots of 1ml from the supernatant were then passed through a 96-well filter (AcroPrep, 1 mL, 1 µm glass fiber) into glass vials, covered with silicone mats and stored at -10 ºC. Chromatographic analyses were conducted on an Agilent 1200 analytical HPLC (high performance liquid chromatography) coupled to an Agilent 6230 Time-of-Flight mass spectrometer via an electrospray ionization source (ESI-TOF; gas temperature: 325 °C, flow: 10 L/m; nebulizer pressure: 35 psig; VCap: 3500 V; fragmentor: 165 V; skimmer: 65 V; octopole: 750 V). Digitoxin (a commercially available cardenolide that has been used as an internal standard in other analyses of saponins (*22*)), internal standard solution (0.50 µL at 0.200 mM MeOH, Sigma-Aldrich), was co-injected with extracts (1.00 µL) and eluted at 0.500 mL/min through a Kinetex EVO C18 column (Phenomenex, 2.1 x 100 mm, 2.6 µ, 100 Å) at 40 °C. Buffers A (water containing 0.1% formic acid) and B (acetonitrile containing 0.1% formic) comprised the linear binary gradient, changing over 30 minutes as follows: 0-1 min 5% B, ramp to 50% B at 4 min, ramp to 100% B at 21 min, 21-25 min 100% B ramping to 1.00 mL/min, before re-equilibrating the column from 25-30 min at 5% B, 0.5 mL/min.

Raw data files were converted to mzML format (*23*) using ProteoWizard MSConvert 3.0 (*24*) before processing using the Bioconductor R package XCMS (*25*). Chromatographic features (retention time and *m/z* bins) were extracted from raw data files before retention time correction using the digitoxin internal standard, peak density grouping and gap-filling. The Bioconductor R package CAMERA (*26*) was then used to identify groups of features (pseudospectra) with similar retention time which were highly correlated across chromatograms ($r > 0.8$), with similar ($r > 0.8$) peak shape and by characteristic isotopic patterns. The feature which was most highly represented across all individual chromatograms was then used as the representative feature from each pseudospectrum. Features were then normalized to plant mass and natural log transformed.

To accommodate differences in ionization efficiency between individual phytochemicals, z transformation was applied to standardize means and variance of all features. Given the large number of plant specimens processed and the considerable amount of HPLC time involved, we applied this correction batch-wise to correct for technical effects arising from changes in instrument response and unavoidable mechanical artifacts such as the changing of the column (the internal standard did not adequately correct for technical error across all compound classes, and was not used for normalization). We applied z transformation across four different batches of samples that were identified through manual inspection of individual compound response across analysis time, and corresponded to analysis batches. In addition, we implemented a "floor" correction across batches so that the highest minimum z score (comparing across batches) became the lowest value for all batches. The latter correction was suggested by the fact that the overall sensitivity of detection varied among batches, with some batches recovering a greater range of small values. Our primary variable of interest (the presence and absence of the butterfly across the landscape) was not confounded with batches (i.e. each batch included samples associated with presence and absence locations). Moreover, we repeated core analyses with different approaches to batch effect correction (e.g., fewer batches or without floor correction) and obtained results that were qualitatively similar to those reported in Fig. 2. In other words, the main result that phytochemical variation has both direct and indirect, negative and positive effects that are comparable in magnitude to other factors (e.g. biotic interactions) is robust to the technicalities of mass spectra processing.

Putative annotations were attempted for all features having factor loadings (see section 5.2 below) with an absolute value greater than 0.3 based on previously described approaches (*8*). Briefly, initial annotation of phenolics (200-400 ppm), saponins (400-650 ppm), lipids and sterols (greater than 400-650 ppm) was done using the relative mass defect (RMD) as a characteristic of each compound (*27*). Annotations were further refined based upon expected retention time, amine-characteristic masses, and molecular ion mass. Features of interest were extracted from raw data and examined for characteristic fragments, adducts and isotopes before cross-referencing against the METLIN mass spectrometric database (*28*) as a way to further categorize annotations. Due to the implications for bioactivity associated with small molecule alkaloids, nitrogenous compounds with ambiguous database hits were characterized as peptides so as to not overstate the presence of defensive alkaloids. One reason we could not annotate potential alkaloids is the lack of literature regarding the presence or characterization of alkaloids in *Medicago,* despite numerous studies of alkaloids in other legumes. Thus, more targeted investigation of *Medicago* alkaloids is warranted. Compounds which did not yield a chemically rational database match as a molecular ion or fragment were classified as unknown.

### 4. Dispersal (estimation of effective migration surface)

We analyzed genotyping-by-sequencing (GBS) data from 541 *Lycaeides melissa* butterflies collected from 27 populations in western North America (see Supplementary Table S2). These DNA sequences were previously described by Chaturvedi et al. (*29*). For the current manuscript, we used the *bwa mem* algorithm (version 0.7.17) (*30*, *31*) to align these data to a new version of the *L. melissa* reference genome (*32*). We ran *bwa mem* with a minimum seed length of 15, considered internal seeds of longer than 20 bp, and only output alignments with a quality score of >30. We then used *samtools* (version 1.5) to compress, sort and index the alignments (*33*). We used *samtools* (version 1.5) and *bcftools* (version 1.6) for variant calling with the original consensus calling algorithm. We used the recommended mapping quality adjustment for Illumina data (-C 50), skipped alignments with mapping quality less than 20, skipped bases with base quality less than 30, and ignored insertion-deletion polymorphisms. We set the prior on SNPs to 0.001 (-P) and called SNPs when the posterior probability that the nucleotide was invariant was ≤0.01 (-p). We filtered the initial set of variants to retain only SNPs with sequence data for at least 80% of the individuals, a mean sequence depth of 2X per individual, at least 10 reads of the alternative allele, a minimum quality score of 30, and no more than 1% of the reads in the reverse orientation (this is an expectation for our GBS method). We then split the SNP data into rare versus common SNPs, as these sets of SNPs can reveal different aspects of demographic history with rare variants being especially informative about recent gene flow and fine-scale population structure (*2*, *34*). We specifically delineated rare variants as those with an overall minor allele frequency of 1-5% (47,470 SNPs) and common variants as those with minor allele frequencies >5% (20,449) (variants with less than 1% frequency were discarded from downstream analyses).

We used *entropy* (version 1.2) to estimate genotypes. This program jointly infers genotypes and allele frequencies while accounting for uncertainty in each, as well as uncertainty in population assignment and ancestry (*2*). The latter is accomplished via an admixture model that assumes the allele copies at each SNP locus are drawn from unknown, hypothetical source populations with each individual having a genome comprised of some mixture of the source populations (*2*). Uncertainty in genotypes comes from limited coverage and sequencing error, as encoded in the genotype likelihoods estimated by *samtools* and *bcftools*. We estimated genotypes assuming two or three source populations. Estimates were obtained via Markov chain Monte Carlo (MCMC) with five chains, each with 5000 iterations as a burn-in followed by 8000 sampling iterations with a thinning interval of 5. Point estimates of genotypes were obtained as the posterior mean estimate of the number of non-reference alleles averaged across chains and numbers of source populations. Genetic distances were then calculated between all pairs of individuals based on average identity by state.

Finally, we estimated relative effective migration rates among populations based on the genetic distances and sampling location; this was done separately for rare versus common variants. Relative effective migration rates were inferred using the program *eems* (version 0.0.0.9000) (*35*). This method does not infer absolute migration rates, but rather identifies regions in space with low or high gene flow relative to a simple two-dimensional stepping-stone isolation-by-distance model (*35*). Thus, this method can identify regions in space receiving limited dispersal or gene flow, even if such regions do not harbor resident butterfly populations (such as, for example, one of our sampled alfalfa locations that has not been colonized by the butterfly). In addition to estimating migration rates separately for rare and common variants, we fit the model (using MCMC with three chains, 4 million sampling iterations, 2 million burnin iterations, and a thinning interval of 10,000) separately assuming 400 demes and 200 demes (to allow for more fine-scale and more coarsely-estimated variation), evenly spaced on a triangular grid.

### 5.1. Analyses: overview

Our ultimate goal was to use structural equation modeling (SEM) to understand potentially complex direct and indirect effects on the presence and absence of our focal butterfly across the landscape. We implemented two levels of data reduction prior to SEM analysis. First, we used exploratory factor analysis on the climate and metabolomic data. Then we used Bayesian ridge regression as a way to reduce the possible number of predictor (exogenous) variables that would need to be included for each response (endogenous) variable in our SEM. Finally, we used the most successful SEM model to discuss the relative importance of different direct and indirect paths affecting presence and absence of the focal butterfly. The success of the SEM model was judged by leave-one-out cross validation and null simulations that permuted the site-level properties (predictor variables) for *L. melissa* presence and absences.

### 5.2. Analyses: factor analysis

Factor analysis is similar to other ordination techniques (such as principal component analysis) in dealing with suites of correlated variables, but the emphasis in factor analysis is on the identification of underlying structure (associated with factors that are not necessarily orthogonal) giving rise to the observed data (*36*). We used the same approach for both datasets (climate and metabolomic), specifically the *factanal* function with promax rotation and Thompson's regression scores. The number of factors calculated for each dataset was determined based partly on inspection of scree plots, but mainly through experimentation by fitting different numbers of factors and repeating downstream analyses (to learn, for example, if additional factors produced additional insight or meaningful effects in SEM models). For the climate dataset, the values going into the factor analysis were already summarized at the site level, thus factor scores could be moved directly into the next analyses. For the metabolomic dataset, data from all 1,651 individual plants were used in the factor analysis, and then average scores at the site level for each factor were retained for ridge regressions and SEM models.

### 5.3. Analyses: Bayesian ridge regression

Before constructing SEMs, we used Bayesian ridge regression as way to focus on a subset of important predictor variables for each of our endogenous variables (presence and absence of the butterfly as well as the abundance of interacting species). Ridge regression is a constrained regression in which coefficients are penalized in a way that reduces coefficients towards zero but does not exclude them (known as an $l_2$ penalty) thus allowing for the simultaneous estimation of effects of a large number of predictors (*37*). In a Bayesian context, ridge regression can be implemented by placing a hyperprior on the precision (= 1/variance) of regression coefficients, which lets the model learn how much the variance on coefficients should be constrained. Ridge regressions were either logistic (for presence and absence of the butterfly) or Gaussian (for logged abundance of ants and functional groups) and were run using JAGS (version 3.2.0) in R with the rjags package (*38*). Minimally-influential priors on regression coefficients were modeled as normal distributions with a mean of zero and precision drawn from a (hyper-prior) gamma distribution (rate = 0.1, shape = 0.1). After sampling with two Markov chains for 100,000 steps each (burnin was not required), performance was evaluated by plotting chain histories, examining effective sample sizes and calculating the Gelman and Rubin convergence diagnostic (*39*, *40*).

All variables were z-transformed prior to analyses, and different subsets of variables were examined for different response variables. For example, for non-herbivorous groups (ants and predators), we allowed for the possibility that plant architecture might be important (as we have seen previously (*7*)), but not phytochemistry, specific leaf area or leaf toughness. Year was included as a binary variable in all ridge regression models to allow for the possibility that variation among the two sampling years should be controlled for while estimating other variables of interest. After ridge regressions had been run (and diagnostics checked) for each response variable, we calculated the fraction of the posterior distribution above or below zero for coefficients whose point estimate (median) was above or below zero, respectively. This value represented our confidence in the sign of the coefficients (positive or negative), and we selected variables (separately for each response variable) with confidence equal to or greater than 75% as variables of potential importance to move forward into SEMs. The cutoff of 75% was determined through experimentation: a higher cutoff missed some variables that were interesting in the downstream SEM, and a lower cutoff included variables that were unimportant (e.g. had very small effect sizes) in the subsequent SEM models.

### 5.4. Analyses: spatial autocorrelation

Our previous work on alfalfa and the Melissa blue butterfly in the Great Basin has suggested to us that spatial autocorrelation might not be a major factor in this system, as spatially proximate patches of alfalfa are in some cases similar and in other cases very different with respect to female oviposition and larval performance (*4*). Nevertheless, the present project encompass more space than our previous ecological work, which raises the importance of quantifying spatial autocorrelation, for which we have taken two approaches. First, we calculated Moran's I statistic of spatial autocorrelation for each of our variables and asked if observations have greater or lesser autocorrelation than would be expected based on 1000 random permutations, using the *moran.randtest* function of the adespatial package (*41*). Moran's I is not, however, appropriate for binary presence and absence data, which is of course the variable of central interest (presence and absence of our focal butterfly). Thus a complementary approach involved the generation of Moran's eigenvector maps (MEMs) which can be used in a multiple regression context to account for spatial autocorrelation at a range of scales (*42*). MEMs were generated using the *dbmem* function in adespatial, and significant MEMs (at $\alpha = 0.05$) were included in a Bayesian logistic ridge regression along with other predictors of Melissa blue presence and absence.

**5.5. Analyses: structural equation models (SEM)**

Following Bayesian ridge regressions we had a suite of predictor variables for each of our four endogenous variables (butterfly presence and absence, ants, tended herbivores, other herbivores, and predators; note that caterpillar parasitoids could have been another endogenous variable in SEM models but were not included as such because they did not emerge as a top variable for the butterfly). Our initial SEM included three unresolved relationships: for example, ants are important for tended herbivores, and (not surprisingly) tended herbivores are important for ants. Unresolved causality can be retained in SEMs as correlations, but inferences are stronger if relationships (directions of influence or association) can be resolved (*43*). To that end, we compared the fit (with AIC) of models with associations pointing in different directions; if the fit in one direction was a dramatic improvement, then the resolved relationship was retained in the final model. We used the piecewise package (*44*) to fit our SEM model, which allows for the inclusion of different error structures (binomial and Gaussian) (*45*). To estimate standardized beta coefficients across the coefficients on different scales, we chose the "Menard.OE" option (*46*), and judged overall model fit using Fisher's C and the associated significance test (*43*). In addition to the traditional $R^2$ reported by the piecewise package, we manually generated a cross validation measure of model fit by repeating the SEM 56 times, leaving out one location with each iteration and calculating the correlation between observed and predicted values for each of our endogenous variables. Finally, we compared the $R^2$ from the full model to the distribution of $R^2$ values from 1,000 null simulations in which site-level attributes were shuffled among locations with each simulation.

For interpretation and visualization, we produced partial plots for certain relationships of interest by reproducing components of the full SEM as stand-alone generalized linear models from which residuals were saved and plotted as either bivariate plots or three-dimensional plots (exploring, for example, the probability of Melissa blue presence or absence as a function of combinations of ant abundance and tended herbivore abundance). In addition, we calculated indirect effects following standard procedures in path analysis involving, for example, the multiplication of coefficients leading from one exogenous variable through an intermediary endogenous variable to a final endogenous variable, as well as the addition of indirect effects to estimate, for example, the total indirect effect associated with a suite of predictor (exogenous) variables of particular type (such as the indirect effect associated with all plant structural exogenous variables) (*43*). We also used direct and indirect path coefficients associated with phytochemical factors as a way to visualize the potential importance of individual compounds. For each compound, we multiplied its loading on a particular factor by the path coefficient associated with that factor, and then summed across factors within indirect effects and direct effects separately.

### Supplementary Results

**1. Arthropod communities, dispersal (effective migration), climate and metabolomics**

Arthropod collections included 20,890 individuals from 1,547 individual plants, and were sorted into 298 individual species (which were morphospecies in the vast majority of cases, in other words identified to taxonomic family and assigned a unique morphospecies number based on phenotype) not including spiders that were not sorted to morphospecies. Family identifications include 123 taxonomic families from 16 orders (summarized in Supplementary Table S3).

Effective migration surfaces for the focal butterfly were estimated for rare and common genetic variants, assuming 200 and 400 demes (surfaces are shown for both classes of variants in the 400 deme model in Supplementary Figure S1). The models were more successful than simple isolation by distance at predicting genetic dissimilarity among spatial units, as can be seen in the comparison in Supplementary Figure S2 for isolation by distance (on the left) vs fitted values on the right based on rare variants in the 400 deme model. Rates of effective migration from all four models (rare and common, 200 and 400 demes) were moved forward into a Bayesian ridge regression predicting Melissa blue presence and absence among locations: the model of rare variants

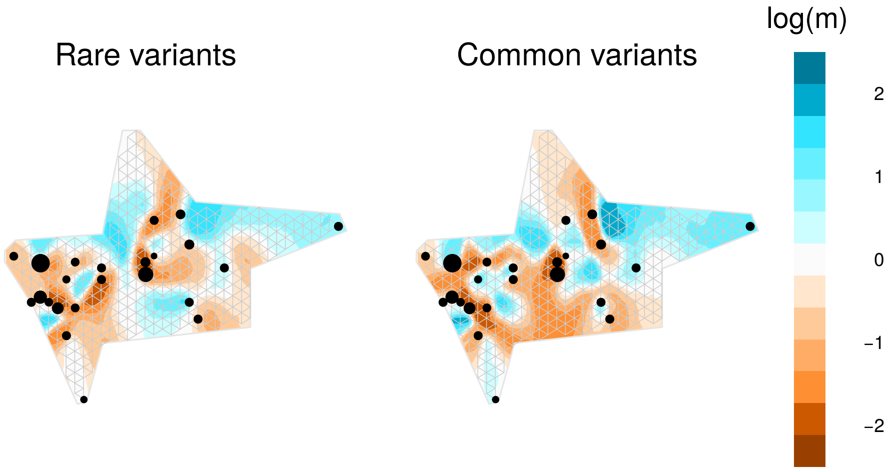


**Supplementary Figure S1.** Relative, effective migration surfaces. Surfaces are shown based on rare (40,449 SNPs) and common (20,449 SNPs) variants. Surface are colored to denote relative migration rates. Points denote demes with circle sizes proportional to sample sizes. The triangular grid used to define deme locations is also shown. In some cases, butterflies from nearby populations could be pooled in a single deme.

at the finer scale of 400 demes was the most influential in the ridge regression (more details below), which is why it is highlighted in Supplementary Figure S2.

Factor analysis was used as a data reduction step before climate and metabolomic data was moved into ridge regressions. We extracted 2 factors for the climate data and 6 for the metabolomic data (Supplementary Figure S3) as optimal in the sense that we maximized the amount of variation captured with the constraint of having few enough factors to be both interpretable and tractable in downstream analyses. In the climate model, the 2 factors explained 72% of the variation among sites, with factor 1 describing a gradient of increasing temperatures and drier conditions, particularly maximum daily temperatures, while factor two is a gradient of increasing minimum temperatures (Supplementary Figure S4). Analysis of the metabolomic data explained 33% of the variation with 6 factors, and we used relative mass defect (RMD) for annotation of major compound classes (Supplementary Table S4 and Supplementary Figure S5). Given the large number of compounds involved, we focused on annotation of the higher-loading compounds as described in Supplementary Methods 3 above.

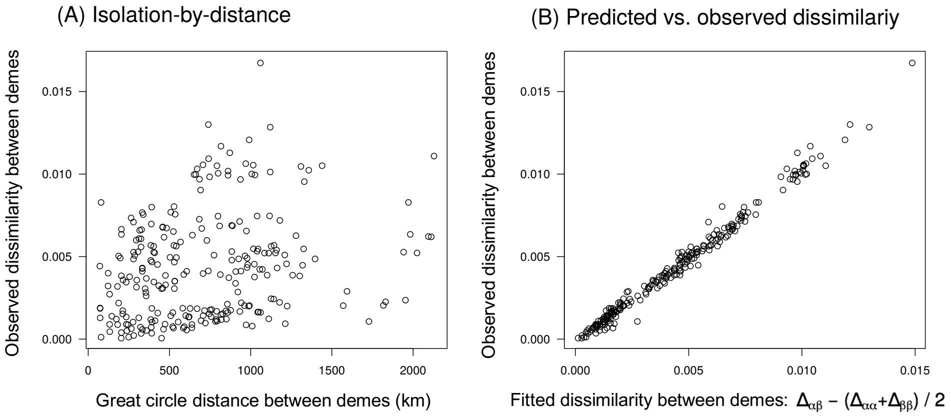


**Supplementary Figure S2.** Observed and expected genetic distances between demes based on a simple isolation-by-distance model (A) and the relative effective migration model (B). In panel (A) great circle distance versus observed genetic dissimilarity is shown with points denoting pairs of demes. In panel (B) the predicted dissimilarity based on the effective migration model versus the observed genetic dissimilarity is shown.

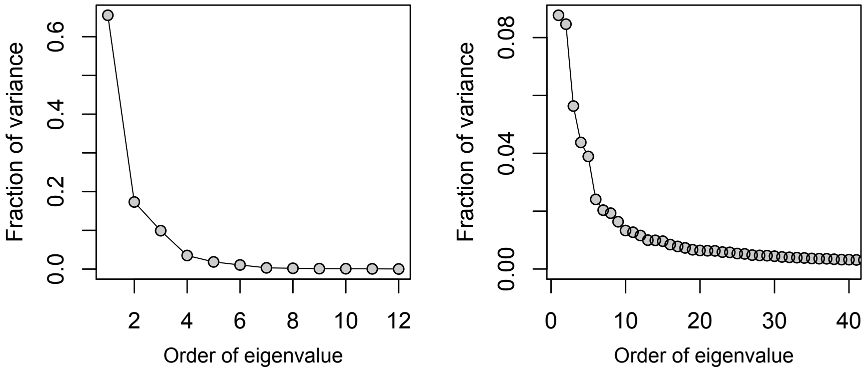


**Supplementary Figure S3.** Scree plots for climate data (on the left) and metabolomic data (on the right). The climate data has 12 variables (12 eigenvalues are shown); the metabolomic data is much larger (849 compounds) so only a portion is shown here.

**2. Ridge regressions**

Bayesian ridge regressions examined five response variables (the focal butterfly, ants, predators, tended herbivores and other herbivores) and a large suite of predictor variables, with slightly different sets of (biologically relevant) predictors in each case. See Supplementary Tables S5 - S9 for the full set of variables considered for each response variable, and Supplementary Figure S6 for pairwise correlations between all variables. All models readily converged and effective sample sizes tended to be in the thousands. Using 75% as the cutoff for confidence in our regression coefficients, we found between 4 and 10 variables that rose to the top as candidates for inclusion in structural equation models (Supplementary Tables S5 - S9). Almost without exception, effects estimated with ridge regressions agreed with *a priori* expectations and previous results in this system. For example, ants had the strongest effect on both Melissa blue presence and absence (*11*) as well as on the abundance of tended herbivores. Specific leaf area (SLA) was highly ranked for the Melissa blue (Supplementary Table S5) with a negative effect as previously observed in an experimental context (*8*), and SLA had a negative effect on other (largely chewing) herbivores (Supplementary Table S8) but not on other ant tended (sucking) herbivores which likely interact differently with physical leaf traits (Supplementary Table S7).

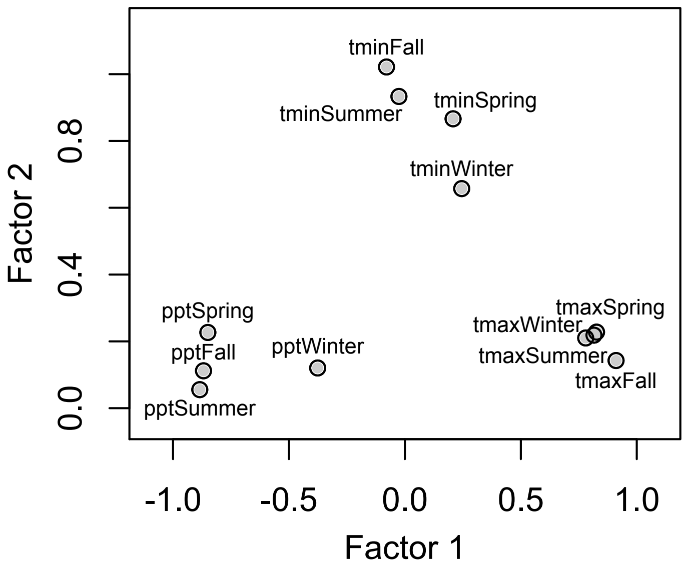


**Supplementary Figure S4.** Climate variable loadings on two axes generated by factor analysis of seasonal values by site for average minimum temperatures ("tmin"), average maximum temperatures ("tmax") and precipitation totals ("ppt").

**3. Spatial autocorrelation**

Spatial autocorrelation was evaluated using Moran's I and comparisons against null simulations. Overall, we found that spatial autocorrelation was low: only a few variables were significantly more clustered on the landscape than would be expected by chance, with (not surprisingly) dispersal having the strongest spatial autocorrelation (Supplementary Table S10). In addition to the tests with Moran's I, we generated MEMs as covariates for spatial structure and included them in a Bayesian ridge regression for our variable of primary interest, the presence and absence of the Melissa blue. Two MEMs fell within the top variables following ridge regression (Supplementary Table S11), using the criterion of 75% confidence. However, comparing Supplementary Table S4 (ridge regression results without MEMs) to Supplementary Table S10 (with MEMs) it can be seen that the vast majority of other variables do not change in their importance while accounting for spatial autocorrelation. Thus we concluded that spatial autocorrelation is present in the system, but do not address it further because it does not alter our primary goal of understanding direct and indirect effects on the presence and absence of the butterfly. Further examination of spatial autocorrelation in this system is in progress in another project examining the relative importance of spatial distance within and among locations for arthropod communities (which can be examined at those two scales, while the presence and absence of the butterfly is only quantified among locations, not within).

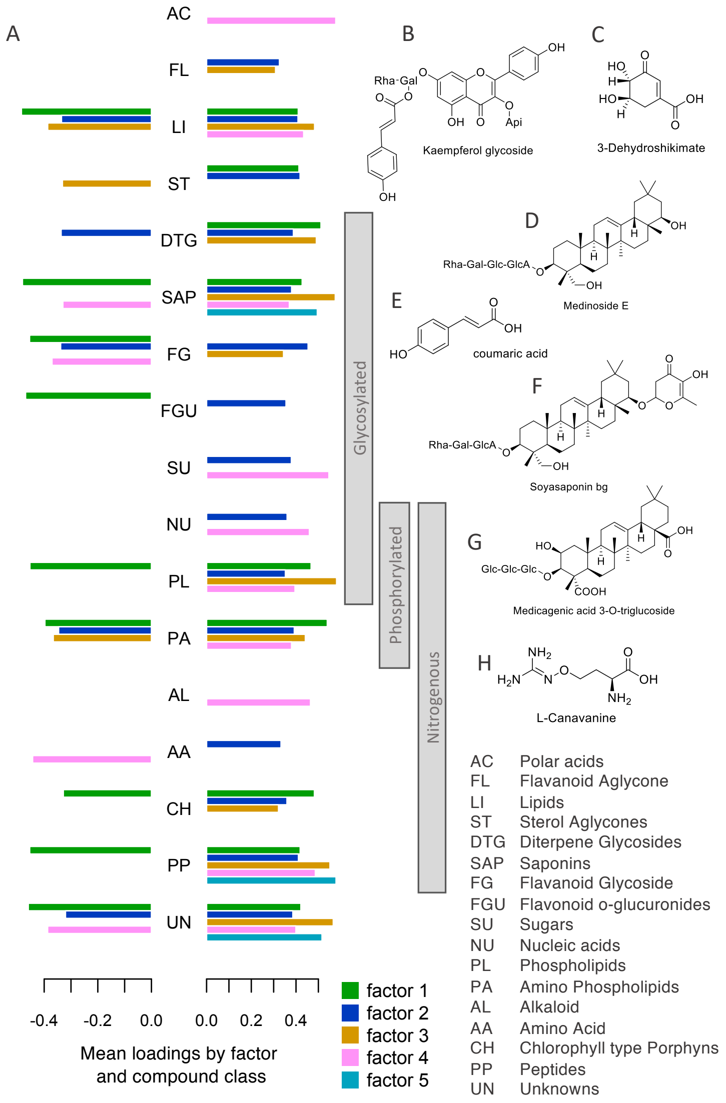


**Supplementary Figure S5.** Summary of factor loadings: horizontal bars (in **A**) show mean loading scores for class assignment groups (see legend key in the lower right). As in Table S4, annotations to compound classes were only determined for features loading at least as high as 0.30 (absolute value) on factors 1 through 5. Vertical gray bars indicate nitrogenous, phosphorylated and glycosylated groups of compound classes. Examples of tentative structural determinations are shown to the right: the compounds in (**B, E and F**) load positively on factor 2 (i.e. have a negative association with tended herbivores); the compounds in (**C** and **H**) load positively on factor 4 (negative association with *L melissa*); and compounds in (**D** and **G**) load positively on both factors 1 and 2.

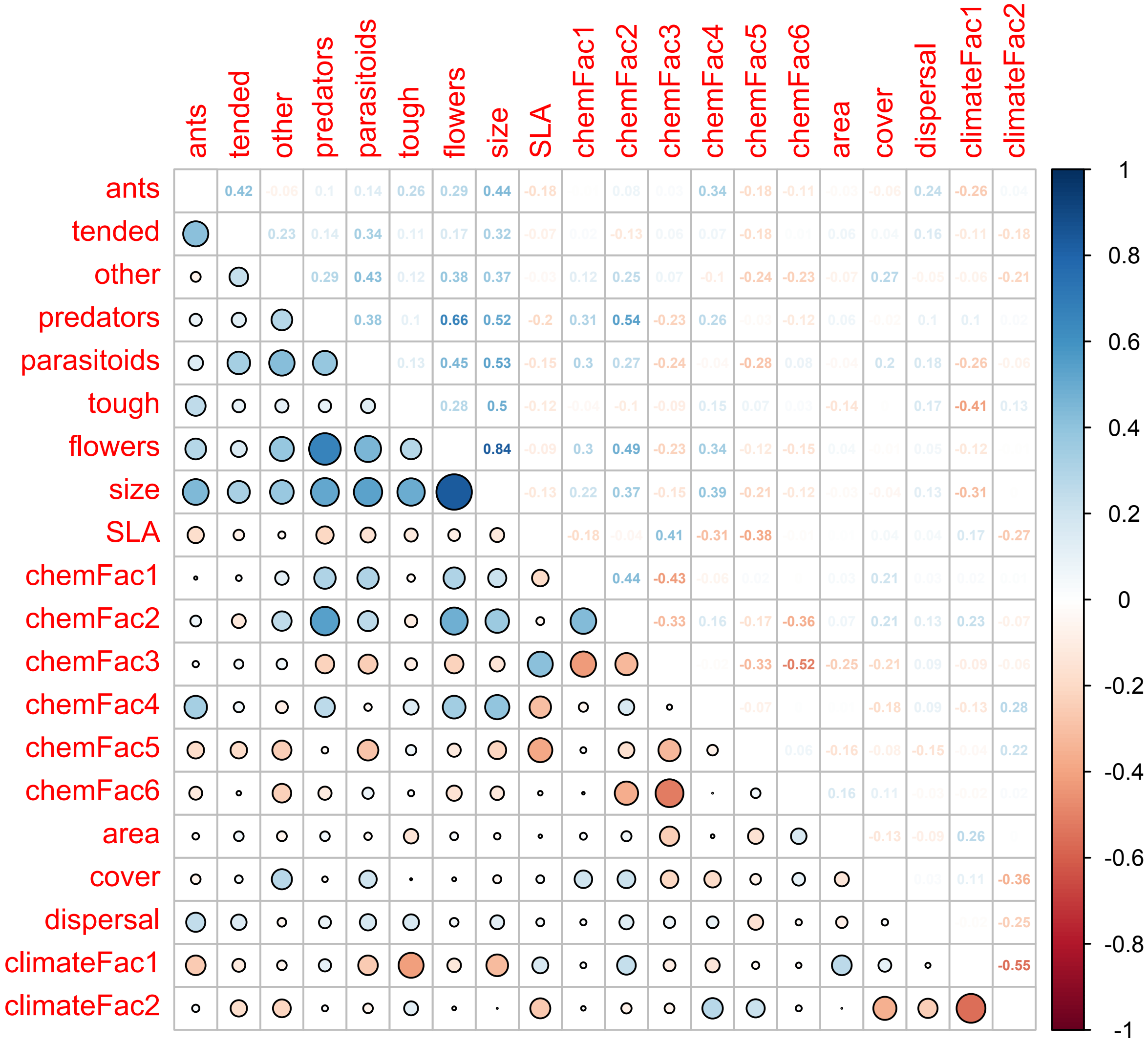


**Supplementary Figure S6.** Pearson correlations coefficients among all endogenous and exogenous response variables. The size of circles in the lower triangle correspond to the magnitude of the correlations (given in the upper triangle, where values closer to zero are faded). The dispersal (effective migration rate) variable shown here is for rare variants at the 400 scale, which was the measure of migration rate that was found to be most relevant to the presence and absence of the focal butterfly (Supplementary Table S4).

### 4. Structural equation models (SEM)

Using the important variables from ridge regressions (Supplementary Tables S5 - S9), we initially constructed a structural equation model that fit the data but that had three unresolved paths (see the base model in Supplementary Table S12). For example, ants are a top variable (in ridge regressions) affecting tended herbivores, and tended herbivores are a top variable for ants, hence the unresolved (or "double headed") path in the base model. Subsequent model comparisons (detailed in Supplementary Table S12) resolved two of those paths, with other herbivores affecting ants and ants pointing to tended herbivores. The latter agrees with our observations: ants (as ecological generalists) can be present without tended herbivores, but tended herbivores are less likely to be successful without ants. The final model is shown in Fig. 2 (with both $R^2$ and leave-one-out correlations) and full results including path coefficients and associated *P* values are given in Supplementary Table S13. In our null model simulations, the variation explained by the real model was roughly three fold greater than the average variation explained in simulated datasets (Supplementary Figure S7).

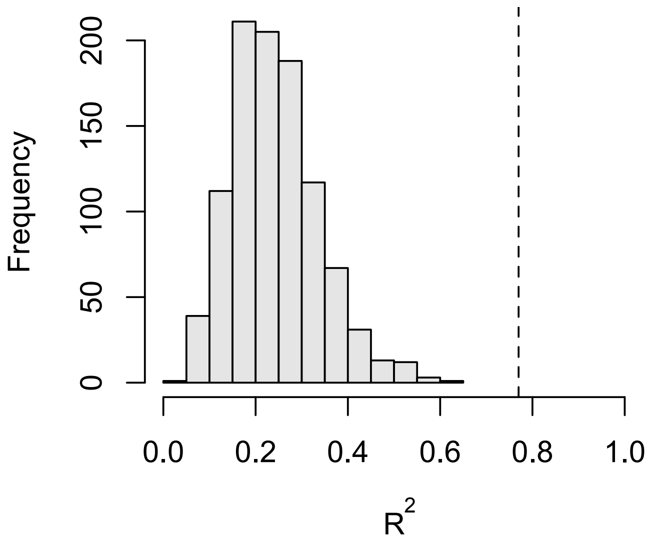


**Supplementary Figure S7.** Histogram of $R^2$ values from 1000 permutations of the data and repeated analysis of the SEM model in Fig. 2. At each permutation, all site-level properties (ant abundance, patch area, specific leaf area, etc.) were randomized among locations. The dotted line shows the actual variance explained from the (non-permuted) model (Fig. 2).

**Supplementary Table S1.** Study locations shown in Fig. 1A; P/A refers to presence or absence of the focal butterfly.

| Code | Population | Lat. | Lon. | P/A |
|---|---|---|---|---|
| 101 | Verdi under the highway | 39.51 | -120.00 | P |
| 102 | Roller Kingdom, Reno, NV | 39.54 | -119.81 | A |
| 103 | Fox Peak Cinema, Fallon, NV | 39.48 | -118.77 | A |
| 104 | Loyalton, CA | 39.67 | -120.23 | A |
| 105 | Rafter 7 Ranch, NV | 38.66 | -118.97 | P |
| 106 | Gardnerville, CA | 38.81 | -119.78 | P |
| 107 | Riverview Dr., NV | 38.91 | -119.72 | A |
| 108 | Lamoille Canyon Rd, NV | 40.72 | -115.49 | P |
| 109 | Lund, NV | 38.88 | -115.02 | A |
| 110 | White Fir St., NV | 39.51 | -119.90 | A |
| 111 | Beckwourth Pass, CA | 39.78 | -120.07 | P |
| 112 | Lupine St, NV | 40.87 | -115.75 | A |
| 113 | Bonneville Shoreline Trl., UT | 41.73 | -111.79 | P |
| 114 | Hardware Ranch, UT | 41.61 | -111.62 | P |
| 115 | South Echo Rd, UT | 41.01 | -111.48 | A |
| 116 | Heber, UT | 40.53 | -111.48 | A |
| 117 | Montague, CA | 41.77 | -122.49 | P |
| 118 | Solar Farm, CA | 41.81 | -121.99 | A |
| 119 | Likely, CA | 41.23 | -120.50 | A |
| 120 | Goose Lake, CA | 41.99 | -120.29 | P |
| 121 | Cedarville, CA | 41.53 | -120.14 | P |
| 122 | Surprise Valley, CA | 41.28 | -120.10 | P |
| 123 | Orovada, NV | 41.57 | -117.80 | A |
| 124 | Winnemucca, NV | 40.98 | -117.74 | A |
| 125 | Girl Farm, CA | 39.63 | -120.00 | P |
| 126 | Verdi Crystal Peak Park, NV | 39.51 | -120.00 | P |
| 127 | Fallon, NV | 39.46 | -118.78 | P |
| 128 | Kingston Canyon (east), NV | 39.21 | -117.12 | P |
| 129 | Star Creek Canyon, NV | 40.55 | -118.12 | P |
| 130 | Kingston Canyon (west), NV | 39.22 | -117.13 | A |
| 131 | Gateway, CO | 38.77 | -108.88 | P |
| 132 | Black Canyon, CO | 38.49 | -107.74 | P |
| 133 | Mainstation Farm, Reno, NV | 39.51 | -119.72 | A |
| 134 | Longley Ln., Reno, NV | 39.47 | -119.77 | A |
| 135 | Quilici Ranch Rd, NV | 39.50 | -119.99 | P |
| 136 | Hemphill Rd., CA | 40.29 | -120.48 | P |
| 137 | Granite Springs, UT | 40.58 | -111.79 | P |
| 138 | Hwy. 189, Heber, UT | 40.49 | -111.42 | A |
| 139 | Hwy. 34, WY | 41.63 | -105.56 | P |
| 140 | Arkansas River, CO | 38.72 | -106.09 | A |
| 141 | Montrose, CO | 38.42 | -107.86 | A |
| 142 | Green River, UT | 39.00 | -110.17 | P |
| 143 | Smith Valley, NV | 38.75 | -119.37 | A |
| 144 | Cokeville, WY | 42.01 | -110.94 | P |

| | | | | |
|---|---|---|---|---|
| 145 | Star Valley, WY | 42.55 | -110.89 | P |
| 146 | Alpine Junction, WY | 43.17 | -111.01 | A |
| 147 | Jackson, WY | 43.48 | -110.77 | A |
| 148 | Victor, ID | 43.64 | -111.11 | P |
| 149 | Elko Campground, MT | 47.92 | -112.76 | A |
| 150 | Moore, ID | 43.73 | -113.37 | A |
| 151 | Richfield, ID | 43.05 | -114.15 | P |
| 152 | Wells, NV | 41.11 | -114.97 | P |
| 153 | Teasdale, UT | 38.31 | -111.49 | P |
| 154 | Helper, UT | 39.65 | -110.86 | P |
| 155 | Tooele, UT | 40.53 | -112.32 | A |
| 156 | Foothill Dr., UT | 40.62 | -111.26 | A |

**Supplementary Table S2.** Populations used for DNA sequencing and estimation of effective migration rates, including the number of individuals (N) sequenced from each location.

| Code | Population | Lat. | Lon. | N |
|---|---|---|---|---|
| ABC | Abel Creek, NV | 41.44 | -117.65 | 19 |
| ABM | Albion Meadows, UT | 40.48 | -111.92 | 46 |
| BHP | Bishop, CA | 37.17 | -118.28 | 20 |
| BSD | Brandon, SD | 43.63 | -96.54 | 20 |
| BST | Bonneville Shoreline Trl., UT | 41.73 | -111.79 | 24 |
| CDY | Cody, WY | 44.51 | -108.98 | 23 |
| CKV | Cokeville, WY | 42.01 | -110.90 | 10 |
| DBQ | De Beque, CO | 39.32 | -108.21 | 20 |
| DCR | Deeth Charleston Rd., NV | 41.30 | -115.38 | 20 |
| GLA | Goose Lake Ag., CA | 41.30 | -120.29 | 20 |
| GVL | Gardnerville, CA | 38.81 | -119.78 | 18 |
| LAN | Lander, WY | 42.65 | -108.36 | 24 |
| LCA | Lam. Canyon, NV | 40.68 | -115.47 | 20 |
| MON | Montrose, CO | 38.37 | -107.82 | 20 |
| MTU | Montague, CA | 41.73 | -122.53 | 19 |
| OCY | Ophir City, NV | 38.94 | -117.24 | 19 |
| REW | Red Earth Way, NV | 38.98 | -118.84 | 20 |
| SCC | Star Creek Canyon, NV | 40.55 | -118.12 | 16 |
| SLA | Silver Lake, NV | 39.66 | -119.93 | 18 |
| SUV | Surprise Valley, CA | 41.28 | -120.10 | 40 |
| TPT | Trout Pond Trailhead, CA | 32.97 | -116.58 | 13 |
| UAL | Upper Alkali Lake, CA | 41.74 | -120.15 | 20 |
| VCP | Verdi Crystal Park, NV | 39.51 | -119.99 | 20 |
| VIC | Victor, ID | 43.66 | -111.11 | 20 |
| WAL | Washoe Lake, NV | 38.65 | -118.82 | 12 |
| YWP | Yellow Pine Camp., WY | 41.25 | -105.40 | 20 |

**Supplementary Table S3.** Summary of arthropod samples by higher taxonomic groups. Count is the total number of individuals for a given taxon, and species is the total number of species (or morphospecies) within a taxon. "NA" indicates families that could not be determined, and also appears in the species richness column for spiders (Araneae) which were not sorted to species (or morphospecies).

| Order (and other higher taxa) | Families | Count | Species |
|---|---|---|---|
| Acari | NA | 113 | 1 |
| Araneae | NA | 1277 | NA |
| Coleoptera | Anthicidae | 30 | 2 |
| Coleoptera | Carabidae | 1 | 1 |
| Coleoptera | Chrysomelidae | 59 | 11 |
| Coleoptera | Cleridae | 1 | 1 |
| Coleoptera | Coccinellidae | 205 | 12 |
| Coleoptera | Coccinellidae | 2 | 1 |
| Coleoptera | Curculionidae | 225 | 7 |
| Coleoptera | Dermestidae | 1 | 1 |
| Coleoptera | Latridiidae | 1 | 1 |
| Coleoptera | Leiodidae | 4 | 1 |
| Coleoptera | Meloidae | 1 | 1 |
| Coleoptera | Melyridae | 172 | 4 |
| Coleoptera | Mordellidae | 7 | 1 |
| Coleoptera | Scraptiidae | 9 | 2 |
| Coleoptera | Staphylinidae | 5 | 2 |
| Coleoptera | Anthicidae | 30 | 2 |
| Coleoptera | Carabidae | 1 | 1 |
| Coleoptera | Chrysomelidae | 59 | 11 |
| Coleoptera | Cleridae | 1 | 1 |
| Coleoptera | Coccinellidae | 205 | 12 |
| Coleoptera | Coccinellidae | 2 | 1 |
| Coleoptera | Curculionidae | 225 | 7 |
| Coleoptera | Dermestidae | 1 | 1 |
| Coleoptera | Latridiidae | 1 | 1 |
| Coleoptera | Leiodidae | 4 | 1 |
| Coleoptera | Meloidae | 1 | 1 |
| Coleoptera | Melyridae | 172 | 4 |
| Coleoptera | Mordellidae | 7 | 1 |
| Coleoptera | Scraptiidae | 9 | 2 |
| Coleoptera | Staphylinidae | 5 | 2 |
| Dermaptera | Forficulidae | 3 | 1 |
| Dermaptera | Forficulidae | 3 | 1 |
| Diptera | Agromyzidae | 2 | 2 |
| Diptera | Anthomyiidae | 5 | 1 |
| Diptera | Asilidae | 2 | 1 |

| | | | |
|---|---|---|---|
| Diptera | Bibionidae | 1 | 1 |
| Diptera | Bombyliidae | 4 | 1 |
| Diptera | Cecidomyiidae | 14 | 2 |
| Diptera | Ceratopogonidae | 144 | 2 |
| Diptera | Chamaemyiidae | 22 | 4 |
| Diptera | Chironomidae | 80 | 2 |
| Diptera | Chloropidae | 93 | 9 |
| Diptera | Chyromyidae | 1 | 1 |
| Diptera | Conopidae | 2 | 1 |
| Diptera | Culicidae | 2 | 1 |
| Diptera | Dolichopodidae | 5 | 1 |
| Diptera | Empididae | 20 | 1 |
| Diptera | Ephydridae | 11 | 3 |
| Diptera | Heleomyzidae | 25 | 2 |
| Diptera | Lauxaniidae | 3 | 2 |
| Diptera | Milichiidae | 4 | 1 |
| Diptera | Muscidae | 1 | 1 |
| Diptera | Mythicomyiidae | 6 | 1 |
| Diptera | Phoridae | 20 | 2 |
| Diptera | Piophilidae | 1 | 1 |
| Diptera | Pipunculidae | 1 | 1 |
| Diptera | Scatopsidae | 5 | 1 |
| Diptera | Scenopinidae | 1 | 1 |
| Diptera | Sciaridae | 20 | 1 |
| Diptera | Sepsidae | 4 | 1 |
| Diptera | Simuliidae | 1 | 1 |
| Diptera | Sphaeroceridae | 1 | 1 |
| Diptera | Stratiomyidae | 2 | 1 |
| Diptera | Syrphidae | 10 | 2 |
| Diptera | Tachinidae | 4 | 3 |
| Diptera | Tephritidae | 17 | 3 |
| Diptera | Agromyzidae | 16 | 1 |
| Diptera | Agromyzidae | 2 | 2 |
| Diptera | Anthomyiidae | 5 | 1 |
| Diptera | Asilidae | 2 | 1 |
| Diptera | Bibionidae | 1 | 1 |
| Diptera | Bombyliidae | 4 | 1 |
| Diptera | Cecidomyiidae | 14 | 2 |
| Diptera | Ceratopogonidae | 144 | 2 |
| Diptera | Chamaemyiidae | 22 | 4 |
| Diptera | Chironomidae | 80 | 2 |
| Diptera | Chloropidae | 93 | 9 |
| Diptera | Chyromyidae | 1 | 1 |
| Diptera | Conopidae | 2 | 1 |
| Diptera | Culicidae | 2 | 1 |
| Diptera | Dolichopodidae | 5 | 1 |
| Diptera | Empididae | 20 | 1 |
| Diptera | Ephydridae | 11 | 3 |
| Diptera | Heleomyzidae | 25 | 2 |

| | | | |
|---|---|---|---|
| Diptera | Lauxaniidae | 3 | 2 |
| Diptera | Milichiidae | 4 | 1 |
| Diptera | Muscidae | 1 | 1 |
| Diptera | Mythicomyiidae | 6 | 1 |
| Diptera | Phoridae | 20 | 2 |
| Diptera | Piophilidae | 1 | 1 |
| Diptera | Pipunculidae | 1 | 1 |
| Diptera | Scatopsidae | 5 | 1 |
| Diptera | Scenopinidae | 1 | 1 |
| Diptera | Sciaridae | 20 | 1 |
| Ephemeroptera | NA | 5 | 1 |
| Hemiptera | Achilidae | 5 | 1 |
| Hemiptera | Anthocoridae | 2019 | 1 |
| Hemiptera | Aphididae | 9266 | 1 |
| Hemiptera | Berytidae | 1 | 1 |
| Hemiptera | Cercopidae | 116 | 3 |
| Hemiptera | Cicadellidae | 1201 | 24 |
| Hemiptera | Cixiidae | 2 | 1 |
| Hemiptera | Clastopteridae | 1 | 1 |
| Hemiptera | NA | 5 | 1 |
| Hemiptera | Coreidae | 30 | 5 |
| Hemiptera | Delphacidae | 92 | 2 |
| Hemiptera | Dictyopharidae | 37 | 1 |
| Hemiptera | Fulgoridae | 4 | 2 |
| Hemiptera | Geocoridae | 401 | 2 |
| Hemiptera | Lygaeidae | 58 | 1 |
| Hemiptera | Membracidae | 902 | 5 |
| Hemiptera | Miridae | 995 | 18 |
| Hemiptera | Nabidae | 108 | 1 |
| Hemiptera | Pentatomidae | 126 | 2 |
| Hemiptera | Psyllidae | 28 | 5 |
| Hemiptera | Reduviidae | 10 | 1 |
| Hemiptera | Rhopalidae | 9 | 1 |
| Hemiptera | Rhyparochromidae | 4 | 2 |
| Hemiptera | Scutelleridae | 9 | 1 |
| Hemiptera | Tingidae | 4 | 1 |
| Hymenoptera | Andrenidae | 2 | 1 |
| Hymenoptera | Aphelinidae | 3 | 1 |
| Hymenoptera | Apidae | 4 | 2 |
| Hymenoptera | NA | 2 | 1 |
| Hymenoptera | Bethylidae | 1 | 1 |
| Hymenoptera | Braconidae | 81 | 9 |
| Hymenoptera | Ceraphronidae | 1 | 1 |
| Hymenoptera | Chalcididae | 2 | 2 |
| Hymenoptera | Chrysididae | 7 | 1 |
| Hymenoptera | Colletidae | 2 | 1 |
| Hymenoptera | Crabronidae | 5 | 3 |
| Hymenoptera | Cynipidae | 2 | 1 |
| Hymenoptera | Dryinidae | 52 | 1 |

| | | | |
|---|---|---|---|
| Hymenoptera | Encyrtidae | 130 | 7 |
| Hymenoptera | Eucharitidae | 2 | 1 |
| Hymenoptera | Eulophidae | 162 | 7 |
| Hymenoptera | Eupelmidae | 11 | 2 |
| Hymenoptera | Eurytomidae | 258 | 3 |
| Hymenoptera | Figitidae | 4 | 2 |
| Hymenoptera | Formicidae | 1510 | 10 |
| Hymenoptera | Halictidae | 10 | 1 |
| Hymenoptera | Ichneumonidae | 6 | 1 |
| Hymenoptera | Mymaridae | 26 | 2 |
| Hymenoptera | Platygastridae | 46 | 7 |
| Hymenoptera | Pompilidae | 1 | 1 |
| Hymenoptera | Pteromalidae | 82 | 5 |
| Hymenoptera | Sphecidae | 1 | 1 |
| Hymenoptera | Torymidae | 61 | 2 |
| Hymenoptera | Trichogrammatidae | 7 | 1 |
| Lepidoptera | Gelechiidae | 16 | 1 |
| Lepidoptera | Geometridae | 5 | 2 |
| Lepidoptera | Hesperiidae | 10 | 2 |
| Lepidoptera | Lycaenidae | 23 | 1 |
| Lepidoptera | NA | 2 | 2 |
| Lepidoptera | Noctuidae | 5 | 1 |
| Lepidoptera | Pieridae | 9 | 1 |
| Lepidoptera | Plutellidae | 1 | 1 |
| Lepidoptera | Pterophoridae | 3 | 1 |
| Lepidoptera | Tortricidae | 1 | 1 |
| Mantodea | Mantidae | 8 | 1 |
| Microcorypia | Meinertellidae | 3 | 1 |
| Neuroptera | Chrysopidae | 10 | 2 |
| Neuroptera | Coniopterygidae | 2 | 1 |
| Neuroptera | Hemerobiidae | 1 | 1 |
| Opiliones | Phalangiidae | 2 | 1 |
| Orthoptera | Acrididae | 181 | 3 |
| Orthoptera | Gryllidae | 15 | 1 |
| Trichoptera | NA | 8 | 1 |

**Supplementary Table S4.** Compound class annotations for all metabolomic features that loaded at least as high as 0.30 on one of the first five factors from exploratory factor analysis (factor 6 was not considered in this context because it had only the most minor association with one of our endogenous variables in SEM). For ease of visualizing which classes load heavily on which factors, loadings less than 0.30 are shown as dashes. For class abbreviations, see Figure S4.

| Feature | Class | Factor 1 | Factor 2 | Factor 3 | Factor 4 | Factor 5 |
|---|---|---|---|---|---|---|
| 2396 | AA | - | 0.3204 | - | - | - |
| 5582 | AA | - | 0.3457 | - | -0.4242 | - |
| 28 | AA | - | - | - | -0.6154 | - |
| 5832 | AA | - | - | - | -0.5154 | - |
| 2722 | AA | - | - | - | -0.4895 | - |
| 5827 | AA | - | - | - | -0.4845 | - |
| 2657 | AA | - | - | - | -0.4839 | - |
| 5923 | AA | - | - | - | -0.3975 | - |
| 5814 | AA | - | - | - | -0.3791 | - |
| 34 | AA | - | - | - | -0.3738 | - |
| 2480 | AA | - | - | - | -0.3729 | - |
| 5826 | AA | - | - | - | -0.3383 | - |
| 2579 | AC | - | - | - | 0.3526 | - |
| 2575 | AC | - | - | - | 0.5909 | - |
| 2580 | AC | - | - | - | 0.6333 | - |
| 2576 | AC | - | - | - | 0.6379 | - |
| 2626 | AC | - | - | - | 0.6754 | - |
| 2572 | AL | - | - | - | 0.3089 | - |
| 2621 | AL | - | - | - | 0.3089 | - |
| 2574 | AL | - | - | - | 0.3968 | - |
| 2570 | AL | - | - | - | 0.4002 | - |
| 2402 | AL | - | - | - | 0.4224 | - |
| 2380 | AL | - | - | - | 0.4494 | - |
| 2726 | AL | - | - | - | 0.5051 | - |
| 2624 | AL | - | - | - | 0.5260 | - |
| 2757 | AL | - | - | - | 0.5380 | - |
| 2573 | AL | - | - | - | 0.5457 | - |
| 2754 | AL | - | - | - | 0.5468 | - |
| 2571 | AL | - | - | - | 0.6310 | - |
| 7716 | CH | -0.3286 | 0.4027 | - | - | - |
| 4720 | CH | 0.3433 | - | - | - | - |
| 5386 | CH | 0.4013 | 0.3067 | - | - | - |
| 7708 | CH | 0.5548 | - | - | - | - |
| 7361 | CH | 0.6295 | - | - | - | - |
| 7713 | CH | - | 0.3210 | - | - | - |
| 64 | CH | - | 0.4078 | - | - | - |
| 60 | CH | - | - | 0.3218 | - | - |

| | | | | | | |
|---|---|---|---|---|---|---|
| 6902 | DTG | 0.5118 | - | - | - | - |
| 6677 | DTG | - | -0.3369 | 0.4910 | - | - |
| 7891 | DTG | - | 0.3403 | - | - | - |
| 47 | DTG | - | 0.3482 | - | - | - |
| 7896 | DTG | - | 0.3657 | - | - | - |
| 7892 | DTG | - | 0.3964 | - | - | - |
| 6904 | DTG | - | 0.4913 | - | - | - |
| 7746 | FL | - | 0.3260 | - | - | - |
| 5692 | FL | - | - | 0.3086 | - | - |
| 6034 | FG | -0.6818 | - | - | - | - |
| 6138 | FG | -0.5307 | - | - | - | - |
| 6221 | FG | -0.5133 | 0.3323 | - | - | - |
| 20 | FG | -0.3617 | 0.5090 | - | - | - |
| 6017 | FG | -0.3212 | - | - | - | - |
| 3362 | FG | -0.3184 | 0.4263 | - | - | - |
| 3239 | FG | - | -0.3386 | 0.3232 | - | - |
| 5882 | FG | - | 0.3211 | - | - | - |
| 8150 | FG | - | 0.3213 | - | - | - |
| 7100 | FG | - | 0.3638 | - | - | - |
| 6022 | FG | - | 0.3982 | - | - | - |
| 6065 | FG | - | 0.4035 | - | - | - |
| 8160 | FG | - | 0.4069 | - | - | - |
| 5861 | FG | - | 0.4099 | - | - | - |
| 5936 | FG | - | 0.4119 | - | - | - |
| 8161 | FG | - | 0.4156 | - | - | - |
| 8162 | FG | - | 0.4163 | - | - | - |
| 5856 | FG | - | 0.4203 | - | -0.3964 | - |
| 35 | FG | - | 0.4323 | - | -0.4152 | - |
| 6269 | FG | - | 0.4383 | - | -0.3470 | - |
| 6063 | FG | - | 0.4445 | - | - | - |
| 6056 | FG | - | 0.4468 | - | - | - |
| 6141 | FG | - | 0.4530 | - | - | - |
| 5999 | FG | - | 0.4552 | - | - | - |
| 6212 | FG | - | 0.4737 | - | - | - |
| 5987 | FG | - | 0.4815 | - | - | - |
| 6014 | FG | - | 0.4825 | - | - | - |
| 8159 | FG | - | 0.4925 | - | - | - |
| 5846 | FG | - | 0.4941 | - | - | - |
| 6071 | FG | - | 0.5197 | - | -0.3707 | - |
| 6067 | FG | - | 0.5211 | - | - | - |
| 6197 | FG | - | 0.5217 | - | - | - |
| 6038 | FG | - | 0.5509 | - | - | - |
| 6037 | FG | - | 0.6462 | - | - | - |
| 5839 | FG | - | 0.6680 | - | - | - |
| 5984 | FG | - | - | 0.3333 | - | - |
| 5990 | FG | - | - | 0.3400 | - | - |
| 6176 | FG | - | - | 0.3805 | - | - |
| 6635 | FG | - | - | - | -0.3250 | - |
| 6091 | FGU | -0.6359 | - | - | - | - |

| | | | | | | |
|---|---|---|---|---|---|---|
| 6027 | FGU | -0.5957 | - | - | - | - |
| 6332 | FGU | -0.5553 | - | - | - | - |
| 95 | FGU | -0.5354 | 0.3237 | - | - | - |
| 6087 | FGU | -0.5270 | - | - | - | - |
| 6079 | FGU | -0.5206 | - | - | - | - |
| 6338 | FGU | -0.4917 | - | - | - | - |
| 6335 | FGU | -0.4845 | 0.3216 | - | - | - |
| 7035 | FGU | -0.4568 | - | - | - | - |
| 5866 | FGU | -0.4307 | - | - | - | - |
| 7046 | FGU | -0.4095 | 0.4243 | - | - | - |
| 7750 | FGU | -0.3730 | - | - | - | - |
| 7563 | FGU | -0.3705 | - | - | - | - |
| 6090 | FGU | -0.3434 | - | - | - | - |
| 6020 | FGU | -0.3110 | 0.3517 | - | - | - |
| 2563 | SU | - | 0.3792 | - | 0.5831 | - |
| 2993 | SU | - | - | - | 0.4034 | - |
| 2541 | SU | - | - | - | 0.4073 | - |
| 2773 | SU | - | - | - | 0.4316 | - |
| 2916 | SU | - | - | - | 0.4402 | - |
| 2721 | SU | - | - | - | 0.4472 | - |
| 2888 | SU | - | - | - | 0.4696 | - |
| 2542 | SU | - | - | - | 0.4915 | - |
| 2712 | SU | - | - | - | 0.5115 | - |
| 2545 | SU | - | - | - | 0.5289 | - |
| 2886 | SU | - | - | - | 0.6074 | - |
| 2543 | SU | - | - | - | 0.6351 | - |
| 2546 | SU | - | - | - | 0.6385 | - |
| 2919 | SU | - | - | - | 0.6567 | - |
| 2765 | SU | - | - | - | 0.6577 | - |
| 2390 | SU | - | - | - | 0.6728 | - |
| 2884 | SU | - | - | - | 0.7224 | - |
| 5535 | LI | -0.6455 | 0.3205 | - | - | - |
| 3678 | LI | -0.5976 | 0.3725 | - | - | - |
| 709 | LI | -0.5640 | 0.3341 | - | - | - |
| 1809 | LI | -0.4872 | 0.4605 | - | - | - |
| 1999 | LI | -0.4012 | 0.4888 | - | 0.3481 | - |
| 5640 | LI | -0.3525 | - | 0.6358 | - | - |
| 82 | LI | -0.3459 | 0.3611 | - | 0.4464 | - |
| 6625 | LI | 0.3039 | - | 0.3802 | - | - |
| 797 | LI | 0.3129 | - | 0.3138 | - | - |
| 5530 | LI | 0.3146 | - | - | - | - |
| 7785 | LI | 0.3320 | - | 0.6140 | - | - |
| 5522 | LI | 0.3332 | - | - | - | - |
| 6951 | LI | 0.3404 | - | 0.3569 | - | - |
| 994 | LI | 0.3433 | 0.4559 | - | - | - |
| 100 | LI | 0.3454 | - | - | - | - |
| 8117 | LI | 0.3556 | - | - | - | - |
| 6942 | LI | 0.3674 | - | -0.3023 | - | - |
| 7960 | LI | 0.3842 | - | - | - | - |

| | | | | | | |
|---|---|---|---|---|---|---|
| 8167 | LI | 0.4092 | - | 0.3899 | - | - |
| 65 | LI | 0.4199 | - | 0.3091 | - | - |
| 6839 | LI | 0.4502 | - | 0.6295 | - | - |
| 728 | LI | 0.4707 | - | - | - | - |
| 8119 | LI | 0.5414 | - | - | - | - |
| 764 | LI | 0.6419 | - | - | - | - |
| 1113 | LI | 0.7163 | - | - | - | - |
| 6436 | LI | - | -0.3355 | 0.5659 | - | - |
| 8011 | LI | - | 0.3320 | - | 0.4391 | - |
| 996 | LI | - | 0.3775 | - | - | - |
| 1806 | LI | - | 0.4021 | - | 0.4909 | - |
| 484 | LI | - | 0.4091 | - | 0.4035 | - |
| 701 | LI | - | 0.4266 | - | - | - |
| 455 | LI | - | 0.4415 | - | 0.4566 | - |
| 7999 | LI | - | 0.4553 | - | 0.4820 | - |
| 136 | LI | - | 0.4980 | - | 0.4089 | - |
| 6950 | LI | - | - | -0.4794 | - | - |
| 6934 | LI | - | - | -0.3859 | - | - |
| 6935 | LI | - | - | -0.3800 | - | - |
| 131 | LI | - | - | 0.3053 | - | - |
| 8165 | LI | - | - | 0.3202 | - | - |
| 175 | LI | - | - | 0.3904 | - | - |
| 7345 | LI | - | - | 0.4144 | - | - |
| 6458 | LI | - | - | 0.4355 | - | - |
| 8135 | LI | - | - | 0.4663 | - | - |
| 3639 | LI | - | - | 0.4745 | - | - |
| 8132 | LI | - | - | 0.4765 | - | - |
| 6433 | LI | - | - | 0.4784 | - | - |
| 5495 | LI | - | - | 0.5916 | - | - |
| 6442 | LI | - | - | 0.6157 | - | - |
| 5666 | LI | - | - | 0.6206 | - | - |
| 6448 | LI | - | - | 0.6572 | - | - |
| 6461 | LI | - | - | 0.6700 | - | - |
| 2470 | NU | - | 0.3302 | - | 0.5467 | - |
| 2469 | NU | - | 0.3903 | - | 0.3417 | - |
| 3141 | NU | - | - | - | 0.3591 | - |
| 2894 | NU | - | - | - | 0.5922 | - |
| 7907 | PA | -0.4836 | 0.4246 | - | - | - |
| 6746 | PA | -0.4523 | 0.4565 | - | - | - |
| 6751 | PA | -0.3494 | - | - | - | - |
| 7367 | PA | -0.3039 | 0.4727 | - | - | - |
| 5806 | PA | 0.3343 | - | -0.4002 | - | - |
| 2326 | PA | 0.3449 | - | - | - | - |
| 5636 | PA | 0.3772 | - | - | - | - |
| 6538 | PA | 0.4198 | - | - | - | - |
| 1112 | PA | 0.4366 | - | 0.4820 | - | - |
| 5672 | PA | 0.4375 | - | 0.5342 | - | - |
| 6570 | PA | 0.4472 | - | - | - | - |
| 4650 | PA | 0.4648 | - | - | - | - |

| | | | | | | |
|---|---|---|---|---|---|---|
| 4474 | PA | 0.4677 | - | 0.5076 | - | - |
| 2291 | PA | 0.4902 | - | - | - | - |
| 1172 | PA | 0.4920 | - | - | - | - |
| 6555 | PA | 0.5004 | - | - | - | - |
| 1111 | PA | 0.5015 | - | 0.4311 | - | - |
| 6054 | PA | 0.5037 | - | - | - | - |
| 3517 | PA | 0.5039 | - | 0.3417 | - | - |
| 7681 | PA | 0.5044 | - | - | - | - |
| 1747 | PA | 0.5191 | - | - | - | - |
| 2322 | PA | 0.5428 | - | - | - | - |
| 5799 | PA | 0.5520 | - | - | - | - |
| 753 | PA | 0.5604 | - | - | - | - |
| 2327 | PA | 0.5737 | -0.3462 | - | - | - |
| 5833 | PA | 0.5949 | - | - | - | - |
| 3686 | PA | 0.5953 | - | - | - | - |
| 7686 | PA | 0.6086 | - | - | - | - |
| 6701 | PA | 0.6090 | - | - | - | - |
| 1130 | PA | 0.6127 | - | 0.3923 | - | - |
| 2275 | PA | 0.6387 | - | 0.3982 | - | - |
| 1207 | PA | 0.6594 | - | - | - | - |
| 6573 | PA | 0.7139 | - | - | - | - |
| 1759 | PA | 0.7379 | - | - | - | - |
| 3672 | PA | 0.7594 | - | - | - | - |
| 6698 | PA | 0.7743 | - | - | - | - |
| 2361 | PA | - | 0.3022 | - | - | - |
| 1249 | PA | - | 0.3043 | - | - | - |
| 2336 | PA | - | 0.3388 | - | 0.3778 | - |
| 5781 | PA | - | 0.3506 | - | - | - |
| 3685 | PA | - | 0.3555 | - | - | - |
| 3679 | PA | - | 0.3567 | - | - | - |
| 1737 | PA | - | 0.3592 | - | - | - |
| 1739 | PA | - | 0.3645 | - | - | - |
| 1209 | PA | - | 0.3677 | - | - | - |
| 1073 | PA | - | 0.3768 | - | - | - |
| 3674 | PA | - | 0.3866 | - | - | - |
| 1721 | PA | - | 0.3940 | - | - | - |
| 1067 | PA | - | 0.4176 | - | 0.3207 | - |
| 774 | PA | - | 0.4907 | - | 0.4146 | - |
| 1066 | PA | - | 0.5434 | - | 0.3309 | - |
| 5768 | PA | - | - | -0.4288 | - | - |
| 4421 | PA | - | - | -0.3852 | 0.3178 | - |
| 6882 | PA | - | - | -0.3569 | - | - |
| 5 | PA | - | - | -0.3546 | - | - |
| 5765 | PA | - | - | -0.3499 | - | - |
| 4452 | PA | - | - | -0.3485 | 0.4149 | - |
| 6883 | PA | - | - | -0.3047 | - | - |
| 5767 | PA | - | - | 0.4473 | - | - |
| 1284 | PA | - | - | - | 0.4844 | - |
| 4890 | PP | -0.7253 | - | - | - | - |

| | | | | | | |
|---|---|---|---|---|---|---|
| 4857 | PP | -0.7082 | - | - | - | 0.3007 |
| 5250 | PP | -0.6735 | - | - | - | 0.3831 |
| 7306 | PP | -0.6310 | - | - | - | 0.3249 |
| 5254 | PP | -0.6176 | - | - | - | 0.3759 |
| 5355 | PP | -0.6101 | - | - | - | 0.3163 |
| 5302 | PP | -0.5806 | - | - | - | 0.3381 |
| 5027 | PP | -0.5340 | 0.3035 | - | - | 0.4859 |
| 5358 | PP | -0.5320 | - | - | - | 0.4307 |
| 4812 | PP | -0.5196 | - | - | - | 0.5438 |
| 5193 | PP | -0.5100 | - | - | - | - |
| 5609 | PP | -0.5069 | - | - | - | - |
| 7148 | PP | -0.4956 | 0.4500 | - | - | - |
| 5208 | PP | -0.4925 | 0.3652 | - | - | - |
| 5330 | PP | -0.4653 | - | - | - | 0.4152 |
| 4903 | PP | -0.4526 | - | - | - | 0.5127 |
| 6702 | PP | -0.4445 | 0.4257 | - | - | - |
| 4949 | PP | -0.4158 | - | - | - | 0.6774 |
| 4798 | PP | -0.4141 | - | - | - | 0.7190 |
| 4917 | PP | -0.4029 | 0.3879 | - | - | - |
| 5301 | PP | -0.3891 | - | - | - | 0.5302 |
| 4931 | PP | -0.3852 | - | - | - | 0.6633 |
| 4845 | PP | -0.3817 | - | - | - | 0.6585 |
| 7438 | PP | -0.3783 | 0.5241 | - | - | - |
| 4979 | PP | -0.3769 | 0.4070 | - | - | - |
| 5054 | PP | -0.3740 | 0.3690 | - | - | - |
| 4978 | PP | -0.3700 | 0.4508 | - | - | - |
| 7446 | PP | -0.3611 | 0.4566 | - | - | - |
| 4848 | PP | -0.3587 | 0.4411 | - | - | - |
| 6364 | PP | -0.3513 | - | - | - | - |
| 5138 | PP | -0.3497 | - | - | - | 0.4059 |
| 5114 | PP | -0.3493 | 0.4151 | - | - | - |
| 5159 | PP | -0.3381 | - | - | - | 0.4317 |
| 4924 | PP | -0.3299 | - | - | - | 0.6657 |
| 4980 | PP | -0.3269 | 0.3881 | - | - | - |
| 5174 | PP | -0.3253 | 0.4781 | - | - | - |
| 4864 | PP | -0.3236 | - | - | - | 0.4736 |
| 4549 | PP | 0.3065 | - | - | - | - |
| 7027 | PP | 0.3080 | - | - | 0.4369 | - |
| 4518 | PP | 0.3141 | - | - | - | - |
| 2486 | PP | 0.3159 | - | - | - | - |
| 5130 | PP | 0.3264 | - | - | - | - |
| 7028 | PP | 0.3349 | - | - | - | - |
| 5223 | PP | 0.3455 | - | - | - | 0.4792 |
| 5292 | PP | 0.3457 | - | - | - | 0.6181 |
| 6118 | PP | 0.3469 | 0.3182 | - | - | - |
| 6116 | PP | 0.3542 | 0.4116 | - | - | - |
| 5143 | PP | 0.3564 | - | - | - | - |
| 6059 | PP | 0.3606 | - | - | - | - |
| 6036 | PP | 0.3622 | - | - | - | - |

| | | | | | | |
|---|---|---|---|---|---|---|
| 7461 | PP | 0.3673 | - | 0.5818 | - | - |
| 4908 | PP | 0.3757 | - | - | - | 0.3651 |
| 5993 | PP | 0.3783 | - | 0.3424 | - | - |
| 5210 | PP | 0.3872 | - | - | - | 0.4326 |
| 5297 | PP | 0.3920 | - | - | - | 0.5847 |
| 5008 | PP | 0.3981 | - | - | - | 0.4246 |
| 5030 | PP | 0.4254 | - | - | - | 0.7083 |
| 36 | PP | 0.4302 | - | - | - | - |
| 4801 | PP | 0.4487 | - | - | - | 0.4264 |
| 3348 | PP | 0.4559 | - | - | 0.3715 | - |
| 5225 | PP | 0.4589 | - | - | - | 0.4508 |
| 6928 | PP | 0.4626 | - | - | - | - |
| 5305 | PP | 0.4863 | - | - | - | 0.5333 |
| 6194 | PP | 0.4870 | - | - | - | - |
| 5379 | PP | 0.5017 | - | - | - | 0.5426 |
| 5253 | PP | 0.5100 | - | - | - | 0.6200 |
| 5137 | PP | 0.5252 | - | - | - | 0.4570 |
| 4958 | PP | 0.5353 | - | - | - | 0.5793 |
| 4820 | PP | 0.5370 | - | - | - | - |
| 5289 | PP | 0.5512 | - | - | - | 0.4254 |
| 4898 | PP | 0.5733 | - | - | - | 0.3530 |
| 5029 | PP | 0.5889 | - | - | - | 0.5647 |
| 4522 | PP | - | 0.3022 | - | 0.3499 | - |
| 5014 | PP | - | 0.3040 | - | - | 0.3124 |
| 6568 | PP | - | 0.3137 | - | - | - |
| 5586 | PP | - | 0.3271 | - | - | - |
| 5590 | PP | - | 0.3277 | - | - | - |
| 5552 | PP | - | 0.3323 | - | - | - |
| 6565 | PP | - | 0.3335 | - | - | - |
| 2548 | PP | - | 0.3362 | - | 0.6040 | - |
| 5612 | PP | - | 0.3407 | - | - | - |
| 7207 | PP | - | 0.3442 | - | - | - |
| 6580 | PP | - | 0.3741 | - | - | - |
| 6556 | PP | - | 0.3874 | - | - | - |
| 6540 | PP | - | 0.4246 | - | - | - |
| 5075 | PP | - | 0.4611 | - | 0.3211 | - |
| 4542 | PP | - | 0.4630 | - | - | - |
| 5556 | PP | - | 0.4691 | - | - | - |
| 4520 | PP | - | 0.4787 | - | - | - |
| 5558 | PP | - | 0.5134 | - | - | - |
| 5580 | PP | - | 0.5288 | - | - | - |
| 29 | PP | - | 0.5335 | - | - | - |
| 4832 | PP | - | 0.5458 | - | 0.3274 | - |
| 5553 | PP | - | 0.5806 | - | - | - |
| 3236 | PP | - | - | 0.3212 | - | - |
| 25 | PP | - | - | 0.7441 | - | - |
| 5787 | PP | - | - | 0.7700 | - | - |
| 5966 | PP | - | - | - | 0.3280 | - |
| 5147 | PP | - | - | - | 0.3389 | 0.3438 |

| | | | | | | |
|---|---|---|---|---|---|---|
| 3212 | PP | - | - | - | 0.3500 | - |
| 6183 | PP | - | - | - | 0.3524 | - |
| 5106 | PP | - | - | - | 0.3629 | 0.4277 |
| 4523 | PP | - | - | - | 0.3950 | - |
| 2646 | PP | - | - | - | 0.4227 | - |
| 2838 | PP | - | - | - | 0.4446 | - |
| 2644 | PP | - | - | - | 0.5758 | - |
| 2547 | PP | - | - | - | 0.5795 | - |
| 2642 | PP | - | - | - | 0.5852 | - |
| 2607 | PP | - | - | - | 0.5902 | - |
| 2638 | PP | - | - | - | 0.5918 | - |
| 2608 | PP | - | - | - | 0.6114 | - |
| 2609 | PP | - | - | - | 0.6232 | - |
| 2611 | PP | - | - | - | 0.6304 | - |
| 2610 | PP | - | - | - | 0.6503 | - |
| 2640 | PP | - | - | - | 0.6528 | - |
| 2634 | PP | - | - | - | 0.6678 | - |
| 7375 | PP | - | - | - | - | 0.3046 |
| 5050 | PP | - | - | - | - | 0.4003 |
| 5012 | PP | - | - | - | - | 0.4361 |
| 5122 | PP | - | - | - | - | 0.4397 |
| 5011 | PP | - | - | - | - | 0.4524 |
| 5113 | PP | - | - | - | - | 0.4707 |
| 5135 | PP | - | - | - | - | 0.4747 |
| 4991 | PP | - | - | - | - | 0.4889 |
| 4874 | PP | - | - | - | - | 0.4927 |
| 5140 | PP | - | - | - | - | 0.4982 |
| 4909 | PP | - | - | - | - | 0.5076 |
| 5049 | PP | - | - | - | - | 0.5205 |
| 5104 | PP | - | - | - | - | 0.5313 |
| 5111 | PP | - | - | - | - | 0.5412 |
| 5046 | PP | - | - | - | - | 0.5916 |
| 5190 | PP | - | - | - | - | 0.5956 |
| 5053 | PP | - | - | - | - | 0.5990 |
| 4974 | PP | - | - | - | - | 0.6010 |
| 4839 | PP | - | - | - | - | 0.6454 |
| 4800 | PP | - | - | - | - | 0.6509 |
| 4816 | PP | - | - | - | - | 0.6578 |
| 4802 | PP | - | - | - | - | 0.6606 |
| 4884 | PP | - | - | - | - | 0.6838 |
| 4902 | PP | - | - | - | - | 0.6988 |
| 5077 | PP | - | - | - | - | 0.7238 |
| 4993 | PP | - | - | - | - | 0.7480 |
| 5081 | PP | - | - | - | - | 0.7484 |
| 4862 | PP | - | - | - | - | 0.7696 |
| 4858 | PP | - | - | - | - | 0.7741 |
| 4954 | PP | - | - | - | - | 0.7788 |
| 4891 | PP | - | - | - | - | 0.8123 |
| 4868 | PP | - | - | - | - | 0.8237 |

| | | | | | | |
|---|---|---|---|---|---|---|
| 4808 | PP | - | - | - | - | 0.8277 |
| 4806 | PP | - | - | - | - | 0.8295 |
| 5037 | PP | - | - | - | - | 0.8500 |
| 4807 | PP | - | - | - | - | 0.8594 |
| 4799 | PP | - | - | - | - | 0.8691 |
| 4815 | PP | - | - | - | - | 0.8860 |
| 5036 | PP | - | - | - | - | 0.8889 |
| 5025 | PP | - | - | - | - | 0.9002 |
| 5023 | PP | - | - | - | - | 0.9023 |
| 5024 | PP | - | - | - | - | 0.9098 |
| 5028 | PP | - | - | - | - | 0.9195 |
| 7992 | PL | -0.7201 | - | - | - | - |
| 4454 | PL | -0.6469 | 0.3051 | - | - | - |
| 195 | PL | -0.6353 | - | - | - | - |
| 196 | PL | -0.5919 | - | - | - | - |
| 780 | PL | -0.5905 | - | - | - | - |
| 7995 | PL | -0.5397 | - | - | - | - |
| 710 | PL | -0.5057 | 0.3147 | 0.3578 | - | - |
| 575 | PL | -0.4782 | - | - | - | - |
| 631 | PL | -0.4669 | - | - | - | - |
| 1276 | PL | -0.4419 | 0.3821 | - | 0.3228 | - |
| 6428 | PL | -0.4361 | - | 0.6815 | - | - |
| 7911 | PL | -0.4211 | - | - | - | - |
| 531 | PL | -0.4017 | - | - | - | - |
| 1297 | PL | -0.3986 | - | - | 0.4148 | - |
| 6454 | PL | -0.3770 | - | - | - | - |
| 4423 | PL | -0.3598 | - | - | - | - |
| 504 | PL | -0.3475 | - | 0.3536 | - | - |
| 6857 | PL | -0.3368 | - | - | - | - |
| 7900 | PL | -0.3263 | - | - | - | - |
| 6487 | PL | -0.3245 | - | - | - | - |
| 75 | PL | -0.3155 | - | 0.5035 | - | - |
| 192 | PL | -0.3088 | - | - | - | - |
| 7679 | PL | 0.3652 | - | - | - | - |
| 4496 | PL | 0.3732 | - | 0.6015 | - | - |
| 8186 | PL | 0.4308 | - | 0.5947 | - | - |
| 1110 | PL | 0.5100 | - | 0.4025 | - | - |
| 2296 | PL | 0.5387 | - | - | - | - |
| 1128 | PL | 0.5878 | - | - | - | - |
| 45 | PL | - | 0.3034 | - | - | - |
| 6771 | PL | - | 0.3076 | - | - | - |
| 57 | PL | - | 0.3077 | - | - | - |
| 4467 | PL | - | 0.3231 | - | 0.6050 | - |
| 516 | PL | - | 0.3290 | - | - | - |
| 6791 | PL | - | 0.3313 | - | - | - |
| 574 | PL | - | 0.3365 | 0.3640 | - | - |
| 7373 | PL | - | 0.3415 | - | - | - |
| 704 | PL | - | 0.3505 | 0.3357 | - | - |
| 6803 | PL | - | 0.3556 | - | - | - |

| | | | | | | |
|---|---|---|---|---|---|---|
| 12 | PL | - | 0.3574 | - | - | - |
| 6747 | PL | - | 0.4081 | - | - | - |
| 773 | PL | - | 0.4360 | - | - | - |
| 691 | PL | - | 0.5030 | - | - | - |
| 601 | PL | - | - | 0.3063 | - | - |
| 649 | PL | - | - | 0.3231 | - | - |
| 794 | PL | - | - | 0.3341 | - | - |
| 1807 | PL | - | - | 0.3378 | 0.3007 | - |
| 626 | PL | - | - | 0.3606 | - | - |
| 62 | PL | - | - | 0.3768 | - | - |
| 1117 | PL | - | - | 0.4138 | - | - |
| 1119 | PL | - | - | 0.4153 | - | - |
| 658 | PL | - | - | 0.4314 | - | - |
| 1698 | PL | - | - | 0.4564 | - | - |
| 625 | PL | - | - | 0.4607 | - | - |
| 7988 | PL | - | - | 0.5542 | - | - |
| 7767 | PL | - | - | 0.5911 | - | - |
| 7787 | PL | - | - | 0.5933 | - | - |
| 2351 | PL | - | - | 0.6155 | - | - |
| 4458 | PL | - | - | 0.6159 | - | - |
| 8181 | PL | - | - | 0.6245 | - | - |
| 7789 | PL | - | - | 0.6296 | - | - |
| 1299 | PL | - | - | 0.6499 | - | - |
| 2087 | PL | - | - | 0.6646 | - | - |
| 7786 | PL | - | - | 0.6990 | - | - |
| 4476 | PL | - | - | 0.7002 | - | - |
| 7790 | PL | - | - | 0.7137 | - | - |
| 4475 | PL | - | - | 0.7393 | - | - |
| 4429 | PL | - | - | 0.7397 | - | - |
| 7924 | PL | - | - | 0.7429 | - | - |
| 7769 | PL | - | - | 0.7439 | - | - |
| 5484 | PL | - | - | 0.7550 | - | - |
| 7989 | PL | - | - | 0.7879 | - | - |
| 68 | PL | - | - | 0.7885 | - | - |
| 4479 | PL | - | - | 0.8165 | - | - |
| 6418 | PL | - | - | 0.8366 | - | - |
| 6431 | PL | - | - | 0.8457 | - | - |
| 4416 | PL | - | - | 0.8460 | - | - |
| 6419 | PL | - | - | 0.8857 | - | - |
| 1995 | PL | - | - | - | 0.3349 | - |
| 3825 | SAP | -0.6596 | 0.3473 | - | - | - |
| 3822 | SAP | -0.5836 | 0.3644 | - | - | - |
| 7519 | SAP | -0.4486 | 0.4072 | - | - | - |
| 6671 | SAP | -0.3715 | - | - | - | - |
| 6670 | SAP | -0.3418 | - | - | - | - |
| 3378 | SAP | 0.3042 | - | - | - | 0.3040 |
| 4927 | SAP | 0.3258 | - | - | - | - |
| 6381 | SAP | 0.3283 | - | 0.5378 | - | - |
| 3338 | SAP | 0.3319 | - | - | - | - |

| | | | | | | |
|---|---|---|---|---|---|---|
| 6623 | SAP | 0.3448 | - | - | - | - |
| 5733 | SAP | 0.3463 | - | 0.6136 | - | - |
| 4925 | SAP | 0.3601 | - | - | - | - |
| 4873 | SAP | 0.3761 | - | - | - | - |
| 7453 | SAP | 0.3808 | - | - | - | - |
| 7405 | SAP | 0.3894 | 0.3245 | - | - | - |
| 4983 | SAP | 0.4014 | - | - | - | - |
| 5756 | SAP | 0.4283 | - | 0.4661 | - | - |
| 5731 | SAP | 0.5293 | - | - | - | - |
| 5722 | SAP | 0.5377 | - | - | - | - |
| 4981 | SAP | 0.5440 | - | - | - | - |
| 4997 | SAP | 0.5621 | - | - | - | - |
| 7509 | SAP | 0.5908 | - | - | - | - |
| 5173 | SAP | 0.6158 | - | - | - | - |
| 7465 | SAP | - | 0.3038 | - | - | - |
| 3321 | SAP | - | 0.3040 | - | - | - |
| 5178 | SAP | - | 0.3064 | - | 0.3042 | - |
| 5013 | SAP | - | 0.3077 | - | - | - |
| 5512 | SAP | - | 0.3119 | - | - | - |
| 7385 | SAP | - | 0.3132 | - | - | - |
| 4985 | SAP | - | 0.3138 | - | - | - |
| 3316 | SAP | - | 0.3222 | - | - | - |
| 3350 | SAP | - | 0.3243 | - | - | - |
| 5719 | SAP | - | 0.3261 | - | - | - |
| 7520 | SAP | - | 0.3261 | - | - | - |
| 7397 | SAP | - | 0.3291 | - | - | - |
| 4964 | SAP | - | 0.3295 | - | - | - |
| 4998 | SAP | - | 0.3405 | - | - | - |
| 4907 | SAP | - | 0.3423 | - | - | - |
| 4926 | SAP | - | 0.3461 | - | 0.3161 | - |
| 3372 | SAP | - | 0.3514 | - | - | - |
| 7422 | SAP | - | 0.3580 | - | - | - |
| 7399 | SAP | - | 0.3609 | - | - | - |
| 5710 | SAP | - | 0.3626 | - | - | - |
| 7249 | SAP | - | 0.3642 | - | - | - |
| 5709 | SAP | - | 0.3670 | - | - | - |
| 7508 | SAP | - | 0.3677 | - | - | - |
| 5167 | SAP | - | 0.3687 | - | - | - |
| 7439 | SAP | - | 0.3705 | - | - | - |
| 5504 | SAP | - | 0.3738 | - | - | - |
| 7423 | SAP | - | 0.3813 | - | - | - |
| 4829 | SAP | - | 0.3814 | - | - | - |
| 7269 | SAP | - | 0.3854 | - | -0.3506 | - |
| 6599 | SAP | - | 0.3940 | - | - | - |
| 7389 | SAP | - | 0.3969 | - | - | - |
| 5072 | SAP | - | 0.3971 | - | - | - |
| 7510 | SAP | - | 0.4010 | - | - | - |
| 7515 | SAP | - | 0.4043 | - | - | - |
| 7418 | SAP | - | 0.4067 | - | -0.3101 | - |

| | | | | | | |
|---|---|---|---|---|---|---|
| 5723 | SAP | - | 0.4379 | - | - | - |
| 7513 | SAP | - | 0.4386 | - | - | - |
| 5168 | SAP | - | 0.4455 | - | - | - |
| 3753 | SAP | - | 0.4473 | - | - | - |
| 7464 | SAP | - | 0.4525 | - | - | - |
| 5207 | SAP | - | 0.4563 | - | - | - |
| 93 | SAP | - | 0.4603 | - | - | - |
| 6634 | SAP | - | 0.4606 | - | - | - |
| 58 | SAP | - | 0.4645 | - | - | - |
| 4937 | SAP | - | 0.4844 | - | - | - |
| 6596 | SAP | - | 0.5151 | - | - | - |
| 7257 | SAP | - | 0.5554 | - | 0.3130 | - |
| 52 | SAP | - | - | 0.3557 | - | - |
| 32 | SAP | - | - | 0.3697 | - | - |
| 6396 | SAP | - | - | 0.4035 | - | - |
| 6380 | SAP | - | - | 0.4790 | - | - |
| 3793 | SAP | - | - | 0.4990 | - | - |
| 5703 | SAP | - | - | 0.5162 | - | - |
| 6371 | SAP | - | - | 0.5174 | - | - |
| 7256 | SAP | - | - | 0.5494 | - | - |
| 5747 | SAP | - | - | 0.5550 | - | - |
| 6627 | SAP | - | - | 0.5551 | - | - |
| 8133 | SAP | - | - | 0.5672 | 0.3482 | - |
| 6391 | SAP | - | - | 0.5701 | - | - |
| 5701 | SAP | - | - | 0.5713 | - | - |
| 6621 | SAP | - | - | 0.5891 | - | - |
| 6641 | SAP | - | - | 0.5941 | - | - |
| 5705 | SAP | - | - | 0.6185 | - | - |
| 6366 | SAP | - | - | 0.6239 | - | - |
| 6638 | SAP | - | - | 0.6268 | - | - |
| 6941 | SAP | - | - | 0.6305 | - | - |
| 3368 | SAP | - | - | 0.6339 | - | - |
| 6369 | SAP | - | - | 0.6414 | - | - |
| 6365 | SAP | - | - | 0.6446 | - | - |
| 6888 | SAP | - | - | 0.6498 | - | - |
| 6367 | SAP | - | - | 0.6598 | - | - |
| 5707 | SAP | - | - | 0.7130 | - | - |
| 6603 | SAP | - | - | 0.7548 | - | - |
| 6880 | SAP | - | - | 0.7633 | - | - |
| 7261 | SAP | - | - | - | 0.4501 | - |
| 7262 | SAP | - | - | - | 0.4901 | - |
| 3308 | SAP | - | - | - | - | 0.3013 |
| 3315 | SAP | - | - | - | - | 0.3015 |
| 3309 | SAP | - | - | - | - | 0.3414 |
| 10 | SAP | - | - | - | - | 0.4162 |
| 4846 | SAP | - | - | - | - | 0.4642 |
| 24 | SAP | - | - | - | - | 0.5304 |
| 4810 | SAP | - | - | - | - | 0.6304 |
| 5156 | SAP | - | - | - | - | 0.8175 |

| | | | | | | |
|---|---|---|---|---|---|---|
| 5154 | SAP | - | - | - | - | 0.8468 |
| 730 | ST | 0.3140 | 0.4141 | -0.3612 | - | - |
| 1715 | ST | 0.3481 | - | - | - | - |
| 2237 | ST | 0.3530 | - | - | - | - |
| 1701 | ST | 0.3777 | - | - | - | - |
| 1714 | ST | 0.5172 | - | - | - | - |
| 2238 | ST | 0.5691 | - | - | - | - |
| 2232 | ST | - | 0.3216 | - | - | - |
| 729 | ST | - | 0.4246 | -0.3026 | - | - |
| 8 | ST | - | 0.4531 | - | - | - |
| 2142 | ST | - | 0.4784 | - | - | - |
| 5727 | UN | -0.6812 | - | - | - | - |
| 5577 | UN | -0.4585 | 0.5042 | - | - | - |
| 2689 | UN | -0.3842 | - | - | - | - |
| 2427 | UN | -0.3132 | - | - | - | - |
| 5160 | UN | 0.3208 | - | - | - | - |
| 6618 | UN | 0.3324 | - | - | - | - |
| 5870 | UN | 0.3452 | - | - | - | - |
| 4987 | UN | 0.3546 | - | - | - | - |
| 5871 | UN | 0.3658 | - | - | - | - |
| 5052 | UN | 0.3747 | - | - | - | 0.3314 |
| 5575 | UN | 0.3810 | 0.3069 | - | 0.3896 | - |
| 5576 | UN | 0.4067 | - | - | - | - |
| 5578 | UN | 0.4228 | - | - | - | - |
| 2582 | UN | 0.5125 | - | - | - | - |
| 6334 | UN | 0.5494 | - | - | - | - |
| 6276 | UN | 0.6967 | - | - | - | - |
| 5132 | UN | - | -0.3205 | - | - | 0.7019 |
| 5635 | UN | - | 0.3236 | - | - | - |
| 974 | UN | - | 0.3259 | - | - | - |
| 7569 | UN | - | 0.4691 | - | 0.3478 | - |
| 5715 | UN | - | - | 0.5059 | -0.3306 | - |
| 6789 | UN | - | - | 0.6275 | - | - |
| 2663 | UN | - | - | - | -0.4952 | - |
| 2674 | UN | - | - | - | -0.3872 | - |
| 2676 | UN | - | - | - | -0.3355 | - |
| 2725 | UN | - | - | - | 0.3035 | - |
| 3242 | UN | - | - | - | 0.4147 | - |
| 2565 | UN | - | - | - | 0.4583 | - |
| 3103 | UN | - | - | - | 0.4842 | - |
| 7305 | UN | - | - | - | - | 0.4670 |
| 55 | UN | - | - | - | - | 0.5654 |

**Supplementary Table S5.** Results from Bayesian ridge regression of variables predicting Melissa blue presence and absence among locations. The last column (probability) shows the fraction of the posterior distribution that is either above or below zero (for positive or negative coefficients, respectively). Variables are sorted by probability; variables with probabilities greater than or equal to 0.75 were used in structural equation models. Credible intervals are 5% equal-tailed probability intervals.

| Variable | Coefficient | Credible interval | Prob. |
|---|---|---|---|
| Ants | 0.88 | (0.12, 2.69) | 0.99 |
| Phytochem. factor 1 | -0.52 | (-1.64, 0.09) | 0.95 |
| Specific leaf area (SLA) | -0.47 | (-1.57, 0.13) | 0.93 |
| Tended herbivores | -0.45 | (-1.61, 0.17) | 0.91 |
| Patch area | 0.40 | (-0.17, 1.38) | 0.90 |
| Predators | -0.36 | (-1.39, 0.28) | 0.86 |
| Phytochem. factor 4 | -0.32 | (-1.49, 0.29) | 0.83 |
| Dispersal (rare, 400) | 0.32 | (-0.43, 1.42) | 0.81 |
| Phytochem. factor 5 | 0.23 | (-0.38, 0.13) | 0.77 |
| Climate factor 2 | 0.22 | (-0.43, 1.2) | 0.75 |
| Leaf toughness | -0.19 | (-1.09, 0.41) | 0.73 |
| Climate factor 1 | -0.18 | (-0.96, 0.49) | 0.72 |
| Parasitoids | -0.16 | (-0.95, 0.53) | 0.69 |
| Dispersal (rare, 200) | 0.18 | (-0.69, 0.17) | 0.68 |
| Dispersal (common, 400) | 0.17 | (-0.71, 0.19) | 0.67 |
| Other herbivores | -0.11 | (-0.81, 0.6) | 0.63 |
| Phytochem. factor 2 | -0.08 | (-0.88, 0.69) | 0.59 |
| Plant cover | -0.06 | (-0.72, 0.62) | 0.58 |
| Phytochem. factor 6 | -0.05 | (-0.76, 0.68) | 0.56 |
| Dispersal (common, 200) | 0.04 | (-0.81, 0.93) | 0.55 |
| Phytochem. factor 3 | -0.04 | (-0.86, 0.76) | 0.55 |
| Floweirng | 0.02 | (-1.09, 0.88) | 0.53 |
| Year | -0.03 | (-0.72, 0.94) | 0.53 |
| Plant size | 0.00 | (-0.81, 0.96) | 0.51 |

**Supplementary Table S6.** Results from Bayesian ridge regression of variables predicting ant abundance among locations. The last column (probability) shows the fraction of the posterior distribution that is either above or below zero (for positive or negative coefficients, respectively). Variables are sorted by probability; variables with probabilities greater than or equal to 0.75 were used in structural equation models. Credible intervals are 5% equal-tailed probability intervals.

| Variable | Coefficient | Credible interval | Prob. |
|---|---|---|---|
| Plant size | 0.35 | (0.08, 0.63) | 0.99 |
| Tended herbivores | 0.28 | (0.05, 0.5) | 0.99 |
| Other herbivores | -0.23 | (-0.48, 0.01) | 0.97 |
| Year | 0.20 | (-0.18, 0.63) | 0.85 |
| Climate factor 1 | -0.08 | (-0.35, 0.21) | 0.71 |
| Predators | -0.06 | (-0.31, 0.19) | 0.68 |
| Plant cover | -0.01 | (-0.23, 0.22) | 0.53 |
| Patch area | 0.00 | (-0.22, 0.22) | 0.51 |
| Climate factor 2 | 0.00 | (-0.26, 0.27) | 0.51 |
| Plant size | 0.35 | (0.08, 0.63) | 0.99 |

**Supplementary Table S7.** Results from Bayesian ridge regression of variables predicting the abundance of ant-tended herbivores among locations. The last column (probability) shows the fraction of the posterior distribution that is either above or below zero (for positive or negative coefficients, respectively). Variables are sorted by probability; variables with probabilities greater than or equal to 0.75 were used in structural equation models. Credible intervals are 5% equal-tailed probability intervals.

| Variable | Coefficient | Credible interval | Prob. |
|---|---|---|---|
| Ants | 0.26 | (0.03, 0.51) | 0.99 |
| Phytochem. factor 2 | -0.21 | (-0.53, 0.06) | 0.94 |
| Other herbivores | 0.13 | (-0.1, 0.37) | 0.86 |
| Plant size | 0.15 | (-0.14, 0.48) | 0.85 |
| Climate factor 2 | -0.12 | (-0.38, 0.12) | 0.84 |
| Patch area | 0.08 | (-0.14, 0.3) | 0.76 |
| Predators | 0.09 | (-0.16, 0.36) | 0.76 |
| Year | 0.11 | (-0.23, 0.45) | 0.74 |
| Leaf toughness | -0.06 | (-0.3, 0.18) | 0.68 |
| Phytochem. factor 5 | -0.05 | (-0.29, 0.19) | 0.66 |
| Phytochem. factor 1 | 0.04 | (-0.19, 0.27) | 0.63 |
| Flowering | -0.04 | (-0.35, 0.24) | 0.60 |
| Climate factor 1 | -0.03 | (-0.29, 0.22) | 0.60 |
| Specific leaf area (SLA) | -0.03 | (-0.27, 0.22) | 0.60 |
| Plant cover | 0.02 | (-0.21, 0.25) | 0.56 |
| Phytochem. factor 3 | 0.02 | (-0.28, 0.3) | 0.56 |
| Phytochem. factor 6 | 0.01 | (-0.25, 0.26) | 0.54 |
| Phytochem. factor 4 | -0.01 | (-0.24, 0.23) | 0.52 |

**Supplementary Table S8.** Results from Bayesian ridge regression of variables predicting the abundance of other herbivores (not ant-tended) among locations. The last column (probability) shows the fraction of the posterior distribution that is either above or below zero (for positive or negative coefficients, respectively). Variables are sorted by probability; variables with probabilities greater than or equal to 0.75 were used in structural equation models. Credible intervals are 5% equal-tailed probability intervals.

| Variable | Coefficient | Credible interval | Prob. |
|---|---|---|---|
| Ants | -0.22 | (-0.47, 0.02) | 0.97 |
| Plant cover | 0.17 | (-0.18, 0.4) | 0.93 |
| Flowering | 0.19 | (-0.32, 0.19) | 0.91 |
| Phytochem. factor 5 | -0.14 | (-0.37, 0.11) | 0.89 |
| Phytochem. factor 4 | -0.14 | (-0.09, 0.35) | 0.88 |
| Tended herbivores | 0.13 | (-0.25, 0.2) | 0.87 |
| Specific leaf area (SLA) | -0.12 | (-0.21, 0.34) | 0.84 |
| Plant size | 0.14 | (-0.19, 0.31) | 0.83 |
| Phytochem. factor 6 | -0.10 | (-0.16, 0.45) | 0.80 |
| Phytochem. factor 3 | 0.10 | (-0.37, 0.09) | 0.77 |
| Year | 0.12 | (-0.22, 0.21) | 0.77 |
| Climate factor 2 | -0.08 | (-0.21, 0.47) | 0.74 |
| Climate factor 1 | -0.06 | (-0.35, 0.15) | 0.69 |
| Predators | 0.06 | (-0.32, 0.17) | 0.69 |
| Phytochem. factor 2 | 0.06 | (-0.38, 0.09) | 0.68 |
| Phytochem. factor 1 | -0.03 | (-0.09, 0.49) | 0.59 |
| Leaf toughness | 0.02 | (-0.21, 0.25) | 0.57 |
| Patch area | 0.00 | (-0.05, 0.39) | 0.52 |

**Supplementary Table S9.** Results from Bayesian ridge regression of variables predicting the abundance of predators among locations. The last column (probability) shows the fraction of the posterior distribution that is either above or below zero (for positive or negative coefficients, respectively). Variables are sorted by probability; variables with probabilities greater than or equal to 0.75 were used in structural equation models. Credible intervals are 5% equal-tailed probability intervals.

| Variable | Coefficient | Credible interval | Prob. |
|---|---|---|---|
| Flowering | 0.51 | (0.21, 0.84) | 0.99 |
| Climate factor 1 | 0.31 | (0.05, 0.58) | 0.99 |
| Climate factor 2 | 0.21 | (-0.04, 0.46) | 0.95 |
| Year | 0.27 | (-0.11, 0.7) | 0.92 |
| Plant size | 0.13 | (-0.2, 0.45) | 0.79 |
| Ants | -0.09 | (-0.32, 0.14) | 0.78 |
| Tended herbivores | 0.06 | (-0.16, 0.27) | 0.70 |
| Other herbivores | 0.04 | (-0.18, 0.27) | 0.65 |
| Plant cover | -0.02 | (-0.22, 0.19) | 0.57 |
| Patch area | 0.00 | (-0.2, 0.2) | 0.50 |

**Supplementary Table S10.** Permutation tests for Moran's I, including the observed value and the probability that the observed value is greater or lesser than the simulated values. Variables are sorted with the strongest autocorrelation at the top. Note that only one dispersal (effective migration rate) variable is shown here as the one that was most important in Bayesian ridge regressions (see Table S5).

| Variable | Observed | *P* |
|---|---|---|
| Dispersal (rare 400) | 0.373 | 0.002 |
| Phytochem. factor 2 | 0.321 | 0.001 |
| Leaf toughness | 0.212 | 0.044 |
| Phytochem. factor 5 | 0.210 | 0.055 |
| Climate factor 1 | 0.196 | 0.057 |
| Parasitoids | 0.190 | 0.063 |
| Climate factor 2 | 0.146 | 0.132 |
| Flowering | 0.092 | 0.287 |
| Plant size | 0.088 | 0.335 |
| Patch area | 0.084 | 0.361 |
| Phytochem. factor 3 | 0.083 | 0.346 |
| Predators | 0.066 | 0.431 |
| Tended herbivores | 0.061 | 0.458 |
| Other herbivores | 0.055 | 0.527 |
| Specific leaf area (SLA) | 0.038 | 0.605 |
| Plant cover | 0.005 | 0.871 |
| Phytochem. factor 4 | -0.023 | 0.941 |
| Phytochem. factor 1 | -0.031 | 0.93 |
| Phytochem. factor 6 | -0.033 | 0.912 |
| Ants | -0.052 | 0.751 |

**Supplementary Table S11.** Results from Bayesian ridge regression of variables predicting Melissa blue presence and absence, as in Table S5, but here including MEMs (covariates for spatial autocorrelation). The last column (probability) shows the fraction of the posterior distribution that is either above or below zero (for positive or negative coefficients, respectively). Variables are sorted by probability; variables with probabilities greater than or equal to 0.75 were used in structural equation models. Credible intervals are 5% equal-tailed probability intervals.

| Variable | Coefficient | Credible interval | Prob. |
|---|---|---|---|
| Ants | 1.33 | (0.19, 25.51) | 0.99 |
| Phytochem. factor 1 | -0.85 | (-16.49, 0.03) | 0.97 |
| MEM7 | 0.89 | (-0.04, 21.01) | 0.97 |
| Tended herbivores | -0.82 | (-19.77, 0.09) | 0.95 |
| Predators | -0.68 | (-12.53, 0.18) | 0.93 |
| Specific leaf area (SLA) | -0.62 | (-12.29, 0.17) | 0.93 |
| Patch area | 0.56 | (-0.19, 8.01) | 0.92 |
| Climate factor 1 | -0.59 | (-14.02, 0.29) | 0.89 |
| Phytochem. factor 4 | -0.55 | (-12.92, 0.26) | 0.89 |
| MEM4 | 0.50 | (-0.35, 7.58) | 0.88 |
| Dispersal (rare, 400) | 0.53 | (-0.47, 13.83) | 0.86 |
| Climate factor 2 | 0.35 | (-0.67, 6.2) | 0.78 |
| Dispersal (rare, 200) | 0.32 | (-1.2, 7.12) | 0.73 |
| Leaf toughness | -0.24 | (-3.54, 0.75) | 0.71 |
| MEM2 | -0.20 | (-3.4, 0.95) | 0.70 |
| Phytochem. factor 3 | -0.21 | (-6.48, 0.99) | 0.68 |
| Parasitoids | -0.16 | (-2.48, 2.15) | 0.64 |
| Dispersal (common, 400) | 0.18 | (-5.84, 2.15) | 0.64 |
| Dispersal (common, 200) | 0.16 | (-2, 4.12) | 0.63 |
| Year | -0.12 | (-9.3, 1.31) | 0.59 |
| Other herbivores | -0.08 | (-1.82, 1.99) | 0.58 |
| Phytochem. factor 6 | -0.07 | (-2.78, 1.42) | 0.57 |
| Plant size | -0.07 | (-4.83, 1.62) | 0.56 |
| Plant cover | -0.06 | (-2.97, 1.44) | 0.56 |
| Phytochem. factor 5 | -0.04 | (-7.18, 0.95) | 0.54 |
| mems.MEM3 | -0.03 | (-3.2, 1.2) | 0.53 |
| mems.MEM1 | 0.02 | (-3.4, 1.49) | 0.52 |
| mems.MEM5 | -0.01 | (-1.1, 2.88) | 0.51 |
| numFlow | 0.00 | (-1.46, 4.34) | 0.50 |
| chemFac2 | 0.00 | (-2.17, 2.61) | 0.50 |

**Supplementary Table S12.** SEM model comparisons addressing unresolved relationships. Shown here is the base model (built from results of Bayesian ridge regressions, see Tables S5-S9) in which unresolved paths appear as "~~", and three sets of pairwise comparisons in which those paths are resolved in different directions, as well as a final model in which the two path resolutions that improved model fit (ants pointing to tended herbivores and other herbivores pointing to ants) are used to modify the base model.

| Models being compared | | AIC |
|---|---|---|
| Base model | | |
| | Melissa = ants + phytochem. 1 + SLA + tended herb. + area + predators + phytochem. 4 + dispersal + phytochem. 5 + climate 2,<br>Other herb. = cover + flowering + phytochem. 5 + phytochem. 4 + SLA + plant size + phytochem. 6 + year + phytochem. 3,<br>Tended herb. = phytochem. 2 + plant size + climate 2 + area + predators,<br>Ants = plant size + year,<br>Predators = flowering + climate 1 + climate 2 + year + plant size + ants,<br>Ants ~~ tended herb.,<br>Ants ~~ other herb.,<br>Tended herb. ~~ other herb. | 157.40 |
| Ants and tended herbivores* | | |
| | With ants as a factor pointing to tended herbivores. | 154.28 |
| | With tended herbivores as a factor pointing to ants. | 156.95 |
| Ants and other herbivores | | |
| | With other herbivores pointing to ants. | 155.84 |
| | With ants pointing to other herbivores. | 158.50 |
| Tended herbivores and other herbivores | | |
| | With other herbivores pointing to tended herbivores. | 158.12 |
| | With tended herbivores pointing to other herbivores. | 159.98 |
| Final model | | |
| | Melissa = ants + phytochem. 1 + SLA + tended herb. + area + predators + phytochem. 4 + dispersal + phytochem. 5 + climate 2,<br>Other herb. = other + cover + flowering + phytochem. 5 + phytochem. 4 + SLA + plant size + phytochem. 6 + year + phytochem. 3,<br>Tended herb. = ants + phytochem. 2 + plant size + climate 2 + area + predators,<br>Ants = plant size + year,<br>Predators = flowering + climate 1 + climate 2 + year + plant size + ants,<br>Tended herb. ~~ other herb. | 153.66 |

** Resolving the relationship between ants and tended herbivores is complicated by the fact that having tended herbivores point to ants creates a loop in which tended herbivores affect ants which affect predators which affect tended herbivores. Thus, to generate the reported comparison, predators were removed as a factor pointing to tended herbivores.*

**Supplementary Table S13.** Path coefficients (including standard errors, standardized coefficients, and *P* values) for the SEM model shown in Fig. 2; the last row reports a correlation coefficient for the unresolved path between tended herbivores and other herbivores.

| Response | Predictor | Estimate | SE | Std. estimate | *P* |
|---|---|---|---|---|---|
| Melissa blue | Ants | 4.31 | 1.42 | 0.63 | 0.002 |
| Melissa blue | Phytochem. fac. 1 | -1.95 | 0.81 | -0.28 | 0.016 |
| Melissa blue | SLA | -1.62 | 0.77 | -0.24 | 0.036 |
| Melissa blue | Tended herb. | -2.26 | 0.82 | -0.33 | 0.006 |
| Melissa blue | Patch area | 1.75 | 0.72 | 0.25 | 0.016 |
| Melissa blue | Predators | -1.22 | 0.65 | -0.18 | 0.058 |
| Melissa blue | Phytochem. fac. 4 | -2.19 | 0.90 | -0.32 | 0.015 |
| Melissa blue | Dispersal (rare 400) | 1.55 | 0.67 | 0.23 | 0.021 |
| Melissa blue | Phytochem. fac. 5 | 1.43 | 0.79 | 0.21 | 0.070 |
| Melissa blue | Climate fac. 2 | 1.09 | 0.65 | 0.16 | 0.094 |
| Other herb. | Plant cover | 0.25 | 0.13 | 0.25 | 0.052 |
| Other herb. | Flowering | 0.39 | 0.23 | 0.39 | 0.092 |
| Other herb. | Phytochem. fac. 5 | -0.19 | 0.14 | -0.19 | 0.171 |
| Other herb. | Phytochem. fac. 4 | -0.30 | 0.14 | -0.30 | 0.037 |
| Other herb. | SLA | -0.24 | 0.16 | -0.24 | 0.143 |
| Other herb. | Plant size | 0.10 | 0.24 | 0.10 | 0.666 |
| Other herb. | Phytochem. fac. 6 | -0.05 | 0.15 | -0.05 | 0.727 |
| Other herb. | Year | 0.18 | 0.28 | 0.09 | 0.523 |
| Other herb. | Phytochem. fac. 3 | 0.24 | 0.19 | 0.24 | 0.219 |
| Tended herb. | Ants | 0.35 | 0.13 | 0.35 | 0.011 |
| Tended herb. | Phytochem. fac. 2 | -0.35 | 0.14 | -0.35 | 0.016 |
| Tended herb. | Plant size | 0.21 | 0.16 | 0.21 | 0.193 |
| Tended herb. | Climate fac. 2 | -0.22 | 0.12 | -0.22 | 0.067 |
| Tended herb. | Patch area | 0.09 | 0.12 | 0.09 | 0.444 |
| Tended herb. | Predators | 0.19 | 0.15 | 0.19 | 0.232 |
| Ants | Other herb. | -0.32 | 0.13 | -0.32 | 0.015 |
| Ants | Plant size | 0.49 | 0.13 | 0.49 | 0.000 |
| Ants | Year | 0.51 | 0.25 | 0.25 | 0.044 |
| Predators | Flowering | 0.66 | 0.18 | 0.66 | 0.001 |
| Predators | Climate fac. 1 | 0.41 | 0.14 | 0.41 | 0.004 |
| Predators | Climate fac. 2 | 0.29 | 0.12 | 0.29 | 0.024 |
| Predators | Year | 0.54 | 0.22 | 0.26 | 0.018 |
| Predators | Plant size | 0.07 | 0.21 | 0.07 | 0.740 |
| Predators | Ants | -0.10 | 0.11 | -0.10 | 0.340 |
| Tended herb. | Other herb. | 0.19 | *NA* | 0.19 | 0.078 |
